# Avl9 defines a family of GTPase-activating proteins that regulate diverse cell biological functions

**DOI:** 10.64898/2026.01.05.697756

**Authors:** Ryan C. Vignogna, J. Christopher Fromme

**Affiliations:** Weill Institute for Cell and Molecular Biology & Department of Molecular Biology and Genetics, Cornell University, Ithaca, New York, USA

## Abstract

Ras-related GTPases are molecular switches regulating hundreds of signaling and trafficking pathways in cells. Many GTPase regulators remain to be identified despite extensive genetic and biochemical screens. Here we present the results of computational protein-protein interaction screens and functional experiments identifying the DENN domain protein Avl9 as a GTPase-activating protein for Arf1. Avl9 is involved in secretion and cell migration, but its molecular function has not been characterized. We determined that Avl9 possesses robust Arf-GAP activity and is recruited to secretory vesicles by Rab8. We find that Avl9 function is conserved in humans and enhances cell migration. We propose that several other DENN domain proteins are also candidate GAPs, and we demonstrate that one candidate previously characterized as a Rab-GEF, DENND6A, exhibits strong Arf-GAP activity towards ARL8B, explaining its role in lysosome positioning. Collectively, this work uncovers a family of monomeric ‘DENN GAP’ proteins that regulate diverse cell biological pathways.

## INTRODUCTION

Ras-related GTPases regulate hundreds of cellular processes by acting as molecular switches to recruit and activate downstream effector proteins^1–5^. The status of the switch is itself regulated by guanine-nucleotide exchange factor (GEF) proteins that turn on the switch by catalyzing GTP-binding, and by GTPase-activating proteins (GAPs) that turn off the switch by stimulating GTP-hydrolysis^6,7^. A central goal in the field is to understand how cells orchestrate the complex network of cellular GTPase pathways by regulating the functions of GEFs and GAPs.

The Arf GTPase family comprises ∼30 proteins in humans, including the endoplasmic-reticulum localized Sar1 and over a dozen different Arf-like (Arl) GTPases, that serve diverse functions at several different organelles^8^. Arf GTPases are particularly important for the regulation of the Golgi complex, recruiting effectors for cargo sorting, vesicle biogenesis and tethering, lipid metabolism and transport, and regulation of other GTPase pathways^9–11^. Arf1, which recruits over a dozen effectors to the Golgi, has also been found to function at other organelles including the plasma membrane^12^, mitochondria^13,14^, and lipid droplets^13,15^. Several GEFs and GAPs for Arf1 at the Golgi have been identified and characterized, but known regulators and effectors of Arf1 beyond the Golgi remain scarce^8^. GEFs and GAPs have been identified through both classical and modern genetic and biochemical screens, but recent studies have demonstrated that protein structure prediction approaches^16,17^ also have great potential to be used for discovery^18–21^.

To identify regulators and effectors of Arf1 that have been missed by other approaches, we employed a structural prediction screen using AlphaFold-multimer^17^. This screen identified a high-confidence prediction between Arf1 and Avl9, a conserved protein of unknown function reported to play roles in secretion and cancer cell migration^22–26^. Avl9 belongs to the DENN domain family of proteins that were originally characterized as Rab-GEFs^27–31^. However, previous work found that Avl9 lacks detectable GEF activity towards an extensive range of Rabs^26,27^. Guided by structural predictions, we used *in vitro* biochemical assays to determine that Avl9 has robust GAP activity towards Arf1. Additional computational screens led to our determination that Avl9 is recruited to secretory vesicles by a direct physical interaction with Rab8. The predicted catalytic residue of Avl9 is important for cancer cell migration in culture, indicating functional conservation and physiological relevance across eukaryotes. We propose that several other DENN domain proteins belong to a monomeric ‘DENN GAP’ family defined by Avl9. We demonstrate that one candidate, human DENND6A, is a GAP for the Arf-family GTPase ARL8B, a finding that elucidates its mechanistic role in positioning lysosomes along the microtubule network in cells.

## RESULTS

### *In silico* screens predict interactions between Avl9 and multiple GTPases

To identify protein-protein interactions involving Arf1 we performed *in silico* protein-protein interaction prediction screens using AlphaFold-multimer^17^. Given the conserved nature of trafficking pathways, we focused on the budding yeast model system due to its relatively small genome. Using Arf1 as a bait, we generated pairwise structural predictions with 450 prey proteins comprising Golgi and post-Golgi vesicle proteomes. We then ranked the resultant bait-prey pairs by their ipTM score, AlphaFold’s measure of confidence in the predicted relative positions of proteins in a complex (**Figure 1A**).

**Figure 1.**
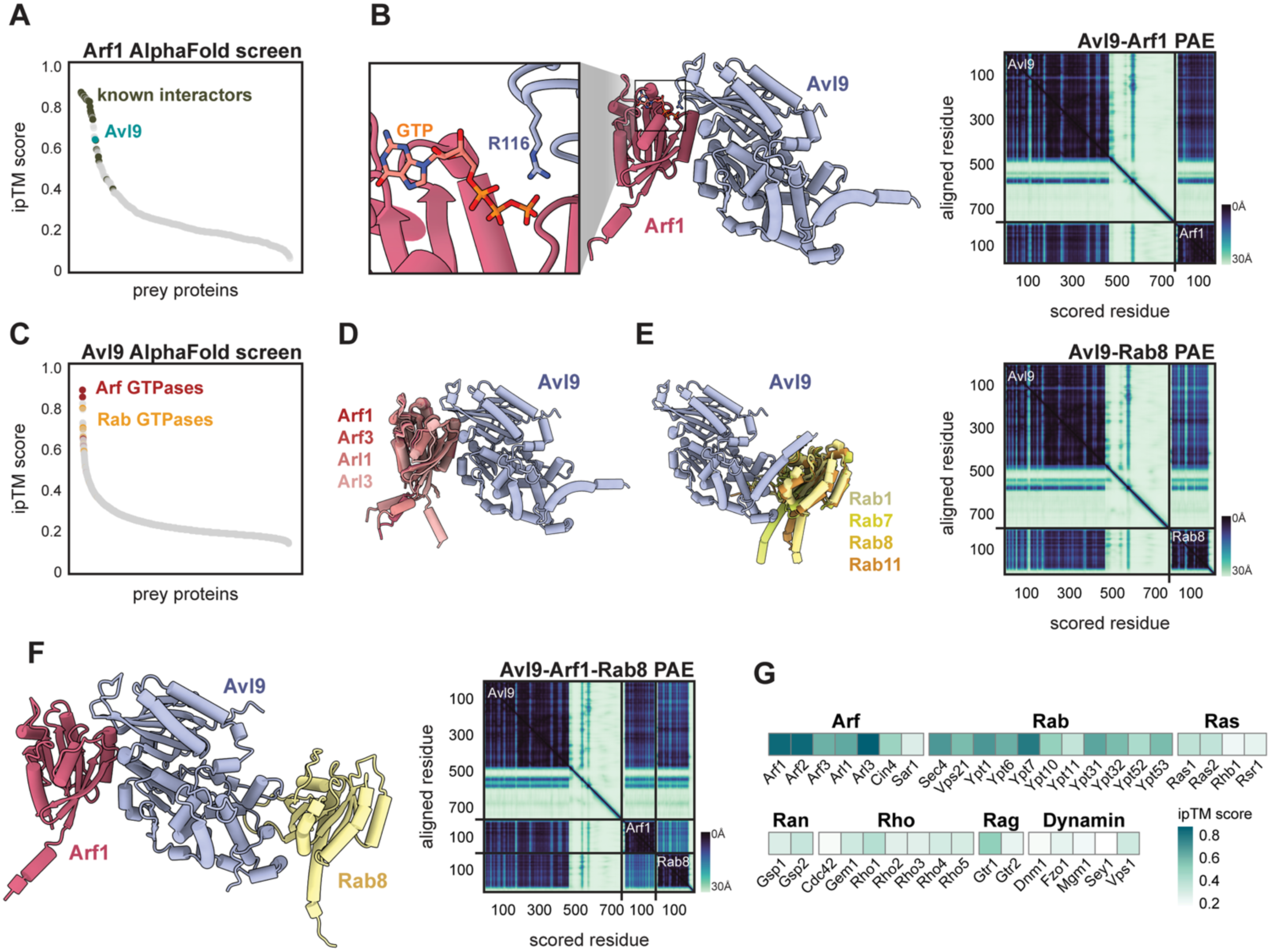
*In silico* screens predict interactions between Avl9 and multiple GTPases. (**A**) Results of a computational protein-protein interaction screen, using AlphaFold-multimer^17^, of Arf1 against proteins involved in Golgi biology. Each point represents the average ipTM score of three structural predictions for a given protein-protein pair. The “known interactors” are experimentally-determined Arf1 physical interactors listed on the *Saccharomyces* genome database^43^. (**B**) *Left,* structural prediction of Arf1 and Avl9, with GTP position overlaid from a crystal structure of Arf1-GTP^36^. The C-terminus of Avl9 is predicted to be disordered and is hidden for clarity. Zoom-in shows the predicted catalytic arginine (R116) of Avl9 pointed towards the terminal phosphate of Arf1-bound GTP. *Right,* predicted alignment error (PAE) plot of Avl9-Arf1 prediction. Scale of expected position error (0Å-30Å) is shown to bottom right of plot. (**C**) Results of a proteome-wide computational interaction screen of Avl9. (**D, E**) Overlaid structural predictions of Avl9 in complex with Arf (left) and Rab (right) GTPases. Representative PAE plot shown. Note the following yeast/human homolog pairs: Arf3/ARF6, Arl3/ARFRP1, Ypt1/Rab1, Ypt7/Rab7, Sec4/Rab8, Ypt31/Rab11 (**F**) Structural prediction of Arf1, Avl9, and Sec4/Rab8 and corresponding PAE plot. (**G**) Heatmap showing the average ipTM score of structural predictions for Avl9 with each of the indicated yeast GTPases.

The screen produced several high-scoring hits, including known Arf1 interactors, such as the COPI vesicle coatomer gamma-subunit Sec21^32^ and the Arf-GEF Sec7^33^. We also found high-confidence predictions involving proteins not previously known to physically interact with Arf1. These include Ent3, which functions with the GGA clathrin adaptor^34^, Sbe2, which is important for yeast cell wall formation^35^, and Avl9, a protein of unknown function that is involved in secretory traffic from the Golgi^22^.

The predicted structure of the Arf1-Avl9 interaction strongly resembles that of a GTPase interacting with a GTPase activating protein (GAP). Superimposing an X-ray crystal structure of GTP-bound Arf1^36,37^ onto the Arf1-Avl9 prediction revealed that R116 of Avl9 is predicted to be positioned in close proximity to the terminal phosphate of GTP, bearing a striking resemblance to the “arginine finger” that many GAP proteins use for GTP hydrolysis^38^ (**Figure 1B**). This prediction was surprising for two reasons: most known Arf-GAPs contain a canonical ArfGAP domain, which Avl9 lacks, and members of the DENN domain protein family to which Avl9 belongs are largely thought to be Rab-GEFs^27,39,40^.

To place the predicted Arf1-Avl9 interaction within the context of a broader set of potential Avl9 interactors, we performed a proteome-wide protein-protein interaction prediction screen with Avl9 as the bait (**Figure 1C**). Among the top hits were Arf1 and several other members of the Arf GTPase family. Superimposing each of these Avl9-Arf predictions indicated that Avl9 was predicted to interact with each Arf GTPase using the same interface (**Figure 1D**). Some GAPs are known to act on multiple closely-related GTPases^41,42^, so it is possible Avl9 acts on multiple Arf substrates, but it is also possible that this screening approach is unable to clearly distinguish between highly similar proteins.

The remaining top-ranked hits of the Avl9 screen included several Rab GTPases (**Figure 1C**). Inspection of these predictions revealed that the Rab GTPases were predicted to interact with a surface of Avl9 that is distinct from that predicted to interact with the Arf GTPases (**Figure 1E**). AlphaFold confidently modeled Avl9 in complex with both an Arf and Rab, indicating these interactions could occur simultaneously (**Figure 1F**). Outside of Arfs and Rabs, no member of other GTPase families were predicted to interact with Avl9 (**Figure 1F**). Based on the results of these *in silico* screens we hypothesized that Avl9 is both an Arf-GAP and a Rab effector.

### Avl9 is an Arf-GAP

To experimentally determine if Avl9 functions as a GAP, we first evaluated the physiological importance of its predicted catalytic arginine residue, which is well conserved among Avl9 homologs (**Figure 2A**). We performed yeast growth assays using a sensitized genetic background in which *AVL9* is essential (*apl2*Δ *vps1*Δ)^22^. We found that the R116A mutant phenocopied *avl9*Δ, without changes in Avl9 expression or localization, indicating this conserved arginine residue is necessary for Avl9 function *in vivo* (**Figures 2B and S1)**.

**Figure 2.**
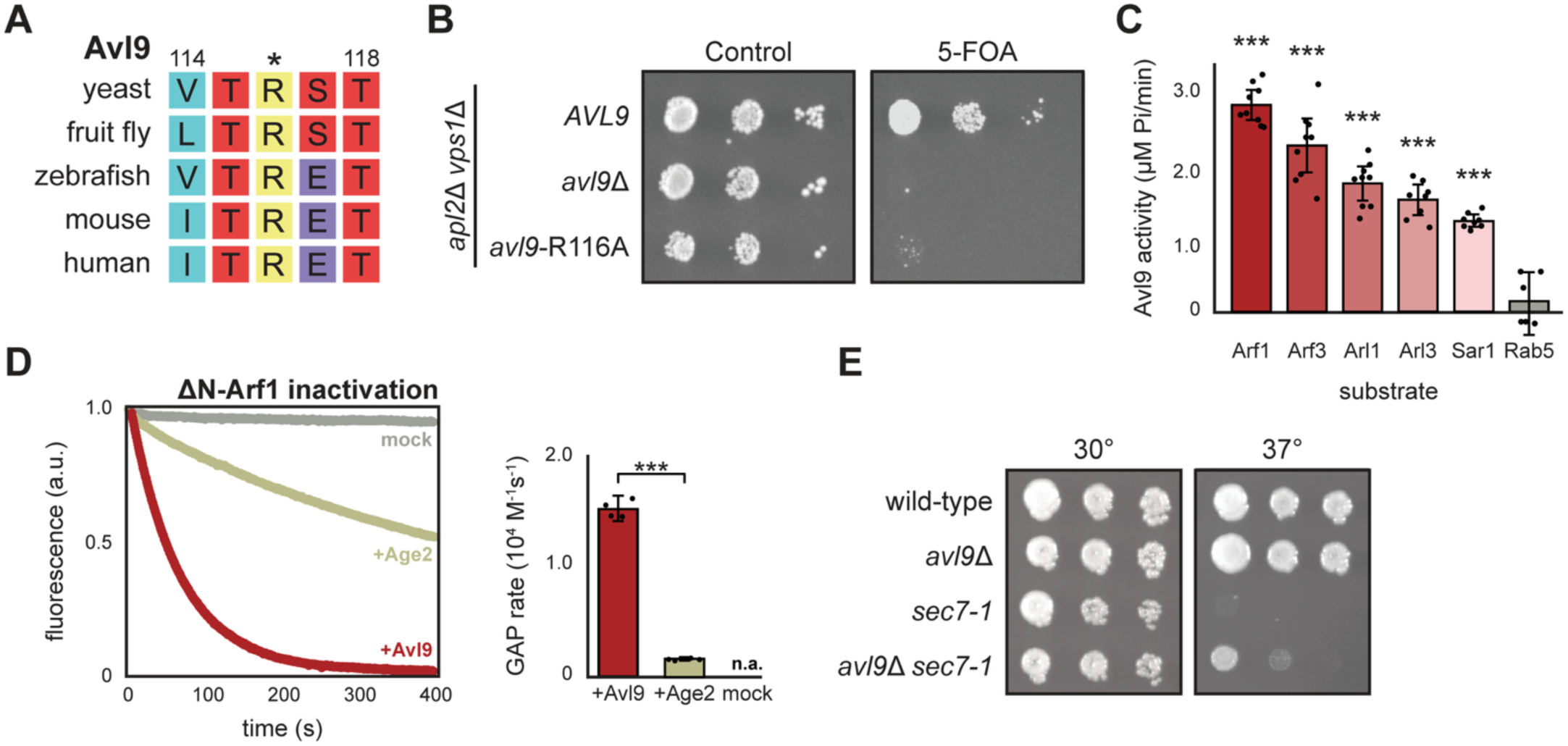
Avl9 is an Arf-GAP. (**A**) Multiple sequence alignment of the arginine finger motif within Avl9 homologs. Numbers indicate residue numbers of the yeast protein and asterisk (*) indicates putative catalytic arginine. (**B**) Yeast “plasmid-shuffling” growth assay, cultures were serially diluted from left to right. The *URA3* covering plasmid encoding *APL2* is counter-selected by 5-FOA. (**C**) Results of malachite green GAP assays with ΔN-Arf substrates. Avl9 activity expressed as the concentration of phosphate (Pi) produced over time. Data points represent technical replicates and error bars represent 95% confidence intervals. Comparisons to Rab5 shown with asterisks. Arf1-Arf3(p=0.0076), Arf1-Arl1(p<0.0001), Arf1-Arl3 (p<0.0001), Arf1-Sar1(p<0.0001), Arf1-Rab5(p<0.0001), Arf3-Arl1(p=0.015), Arf3-Arl3(p=0.0002), Arf3-Sar1(p<0.0001), Arf3-Rab5(p<0.0001), Arl1-Arl3(p=0.71), Arl1-Sar1(p=0.023), Arl1-Rab5 (p<0.0001), Arl3-Sar1(p=0.47), Arl3-Rab5(p<0.0001), Sar1-Rab5(p<0.0001) (df=5, *F*=61, one-way ANOVA with Tukey post hoc). (**D**) Intrinsic tryptophan fluorescence GAP assay with ΔN-Arf1. *Left*, representative fluorescence traces for reactions with Avl9, Age2, or mock (a.u. = arbitrary units). *Right*, quantification of GAP rates; no rate could be assigned to mock reactions (n.a.). Data points represent rates from replicate experiments and error bars represent 95% confidence intervals. Comparison of Avl9 and Age2 GAP rates shown with asterisks (t=35.1, df=3.11, p<0.0001, Welch’s modified t-test). (**E**) Yeast growth assays, cultures were serially diluted from left to right on YPD and grown at the indicated temperature.

To directly test GAP activity and substrate specificity *in vitro* we purified Avl9 and soluble versions of the five yeast Arf GTPases involved in membrane trafficking (**Figure S2A**). We used these reagents to perform biochemical GAP assays quantifying the concentration of free phosphate produced from hydrolysis of GTP bound to the GTPases. Avl9 exhibited significant GAP activity towards all five Arf proteins, with the highest activity towards Arf1, but was not active on Rab5 (yeast Vps21) (**Figure 2C**). To measure the kinetic rate of Avl9 GAP activity, we used an established assay that monitors the intrinsic tryptophan fluorescence of Arf1 as a readout of its activation state in real time^44,45^. Under these conditions, Avl9 exhibited robust GAP activity towards Arf1, with a rate of 1.5 x 10^4^ M^-1^s^-1^, which was tenfold higher than that of the established Arf-GAP Age2 (SMAP2 homolog) (**Figures 2D and S2B**)^44^. These results indicate that Avl9 is a potent Arf-GAP capable of inactivating multiple Arf GTPases *in vitro*.

As observed previously in large-scale imaging screens^46–48^, Avl9 localizes to the bud neck and bud tip of yeast cells (**Figure S1**), indicating that Avl9 resides on secretory vesicles (which we confirm by colocalization below). While Avl9 is able to inactivate multiple Arf GTPase substrates *in vitro*, its localization to secretory vesicles likely constrains its specificity *in vivo*. Arf-GAPs act as GTPase ‘erasers’, ensuring that Arf GTPases are removed from compartments where they do not belong. We hypothesize that Avl9 may function to ensure that no Arf GTPases localize to secretory vesicles. One expectation is that substrate GTPases may localize ectopically in cells lacking their GAPs. We therefore examined the localization of Arf GTPases in *avl9*Δ cells, but none were observed to mislocalize. Given the promiscuity of GAPs, it appears likely that other Arf-GAPs, such as Gcs1, Age2, Glo3, and Gts1, provide redundant Arf-GAP function in the absence of Avl9. This redundancy is consistent with the fact that each of these proteins are individually non-essential^49–51^. We therefore sought an alternative method to identify a physiological role for Avl9.

Arf1 was the best Avl9 substrate in vitro and, given its role in vesicle formation at the trans-Golgi network^52,53^, is a strong candidate Avl9 substrate on secretory vesicles. We therefore wondered whether *AVL9* exhibits a positive genetic relationship with *SEC7*, the GEF (i.e. activator) for Arf1 during secretory vesicle formation^33^. At high temperature, cells harboring the temperature-sensitive *sec7-1* allele were inviable, but this growth defect was partially rescued in the *sec7-1 avl9*Δ double mutant strain (**Figure 2E**). The small size of the effect is consistent with the fact that Sec7 activates Arf1 to drive several different Golgi trafficking pathways, whereas Avl9 appears to be specific for secretory traffic^22^. Thus, phenotypic suppression of an Arf-GEF mutant by loss of Avl9 provides evidence that Avl9 negatively regulates Arf1 *in vivo*.

### Avl9 is recruited to secretory vesicles by Rab8

Our computational screens resulted in high-confidence predictions of interactions between Avl9 and Rab GTPases, involving a surface distinct from the catalytic site of Avl9 (**Figure 1F**). Based on these predictions we hypothesized that a Rab protein mediates Avl9 localization to secretory vesicles. To precisely determine the localization of Avl9 and assess potential Rab interaction partners, we measured colocalization of Avl9 with Rab8 (yeast Sec4) and Rab11 (yeast Ypt31). Rab11 is present both at the Golgi and secretory vesicles while Rab8 is the principal secretory vesicle-associated Rab^54^. We found that Avl9 colocalizes very well with Rab8 but only occasionally with Rab11 (**Figures 3A and 3B**). The small amount of colocalization with Rab11 occurred on secretory vesicles at the bud neck and tips of growing buds, and not at the Golgi (**Figure 3C**). Taken together these results establish that Avl9 colocalizes strongly with Rab8 on secretory vesicles.

**Figure 3.**
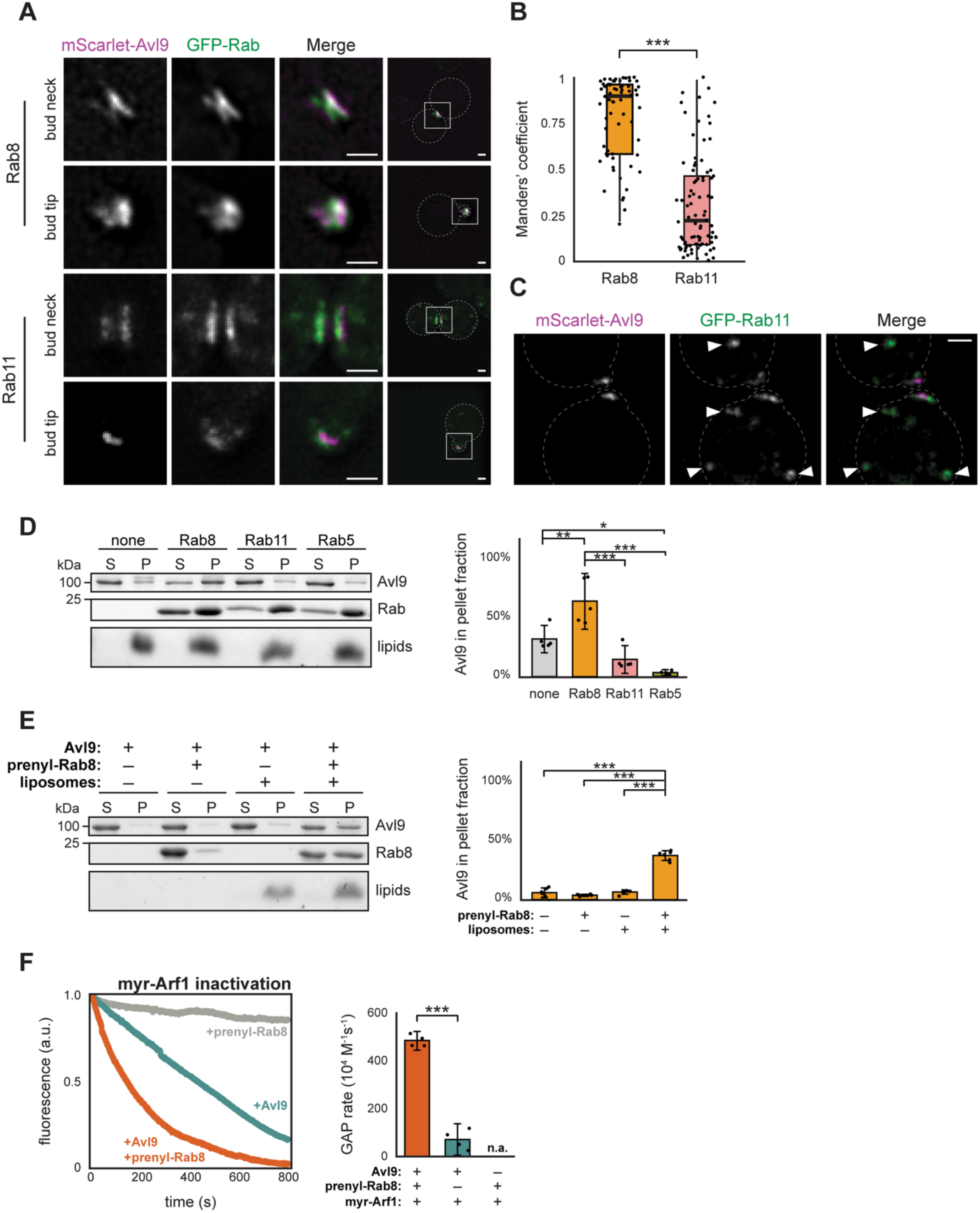
Rab8 colocalizes with Avl9, recruits it to membranes, and stimulates its Arf-GAP activity. (**A**) Live cell fluorescence microscopy images. Scale bars represent 1 μm. (**B**) Quantification of GFP overlap with respect to mScarlet, based on images of entire yeast cells. Data points from individual images shown as dots. Bold lines represent data medians, boxes extend to the first and third quartile of data, and whiskers extend to furthest data point within 1.5x of the interquartile range of the box. Comparison of mean Manders’ coefficient for Rab8 (*n*=65) and Rab11 (*n*=85) groups shown (t=11.8, df=142.9, p<0.0001, Welch’s modified t-test). (**C**) Live cell fluorescence microscopy images. Scale bars represent 1 μm. Arrowheads point to Golgi puncta. (**D**) Representative gel (left) and quantification (right) of pelleting assays with Avl9, His-tagged Rabs, and Ni^2+^ Golgi liposomes. S=supernatant fraction and P=pellet (membrane-bound) fraction. Data points represent measurements from replicate experiments and error bars represent 95% confidence intervals. None-Rab8(p=0.0037), none-Rab11(p=0.14), none-Rab5(p=0.014), Rab8-Rab11(p<0.0001), Rab8-Rab5(p<0.0001), Rab11-Rab5(p=0.52) (df=3, *F*=22, one-way ANOVA with Tukey post hoc). (**E**) Representative gel (left) and quantification (right) of Avl9 pelleting assays with or without prenylated Rab8 and Golgi liposomes. Data points represent measurements from technical replicates and error bars represent 95% confidence intervals. Comparisons of +prenyl-Rab8/+liposomes versus –prenyl-Rab8/–liposomes (t=14.0, df=10.0, p<0.0001), +prenyl-Rab8/–liposomes (t=20.5, df=5.37, p<0.0001), –prenyl-Rab8/+liposomes (t=17.7, df=6.85, p<0.0001) (Welch’s modified t-test) (**F**) Native tryptophan fluorescence GAP assay with myristoylated Arf1. *Left,* representative fluorescence traces for reactions with or without Avl9 or prenyl-Rab8. *Right*, quantification of GAP; no rate could be assigned to reactions where Avl9 was excluded (n.a.). Data points represent rates from replicate experiments and error bars represent 95% confidence intervals. Comparison of +Rab8/+Avl9 and –Rab8/+Avl9 shown (t=17.2, df=4.89, p<0.0001, Welch’s modified t-test).

We performed *in vitro* membrane-binding assays to test whether Avl9 directly interacts with Rab11 or Rab8. We first performed these experiments with liposomes containing 5% Ni^2+^-DOGS lipid to tether purified His-tagged Rab proteins to the membrane surface. We found that while Avl9 exhibited some intrinsic binding to the liposomes, significant additional recruitment to liposomes was induced by Rab8, but not by Rab11 nor Rab5 (**Figure 3D**). This result is concordant with the *in vivo* colocalization and together these results indicate Rab8 is the most likely Rab binding partner of Avl9.

Native Arf proteins bind to lipid membranes when they are GTP-bound. We therefore tested the effect of Rab8 on Avl9 Arf-GAP activity using an established physiological *in vitro* GAP assay in which purified, full-length myristoylated Arf1 (myr-Arf1) is bound to synthetic liposome membranes^44,45^. First, we confirmed that GTP-bound Rab8, when physiologically anchored to liposomes via prenylation of its C-terminus (prenyl-Rab8), recruits Avl9 to liposome membranes (**Figures 3E and S3A**). We then measured Avl9 Arf-GAP activity towards liposome-bound myr-Arf1 with or without addition of prenyl-Rab8. In the absence of prenyl-Rab8, the GAP rate of Avl9 (69 x 10^4^ M^-1^s^-1^) was 45-fold higher towards myr-Arf1 compared to ΔN-Arf1, likely due to increased encounters between Avl9 and myr-Arf1 on liposomes (**Figures 3E and S3B**). When prenyl-Rab8 was bound to liposomes together with myr-Arf1, the GAP rate of Avl9 (480 x 10^4^ M^-1^s^-1^) was an additional 7-fold higher (**Figure 3F**). Therefore, Rab8 recruitment of Avl9 to membranes significantly enhances its Arf1-GAP activity. We note that this reaction represents an example of GTPase crosstalk in which the active form of one GTPase (Rab8) directs the inactivation of another GTPase (Arf1).

To assess the importance of the Rab8-Avl9 interaction for localizing Avl9 *in vivo*, we used the predicted structure to design mutations expected to disrupt this interaction. We observed that Rab-Avl9 predictions generated by AlphaFold3^55^ yielded different results than those produced by AlphaFold2, and suggested an additional potential Rab binding site on Avl9 (referred to as ‘site B’) (**Figures 4A, S4A, and S4B**). We therefore introduced mutations into both of the predicted Rab-binding interfaces of Avl9 and observed localization of the mutant constructs *in vivo*. Mutations in either interface caused Avl9 to become significantly mislocalized and dispersed throughout the cytoplasm (**Figure 4B**) without affecting protein levels (**Figure 4C**), indicating that both sites are important for Avl9 localization to secretory vesicles. Interestingly, mutations in Rab site A are viable in the sensitized *apl2*Δ *vps1*Δ background whereas mutations in Rab site B are inviable in this background (**Figure S3C**). One interpretation of these results is that the site A mutant protein retains a low level of normal localization that is sufficient to provide function, whereas the site B mutant protein does not. Although it is possible that Avl9 uses one or both of these conserved surfaces to bind to some other recruiting factor, the *in vitro* and *in vivo* results provide strong support for the hypothesis that Avl9 is a Rab8 effector on secretory vesicles.

**Figure 4.**
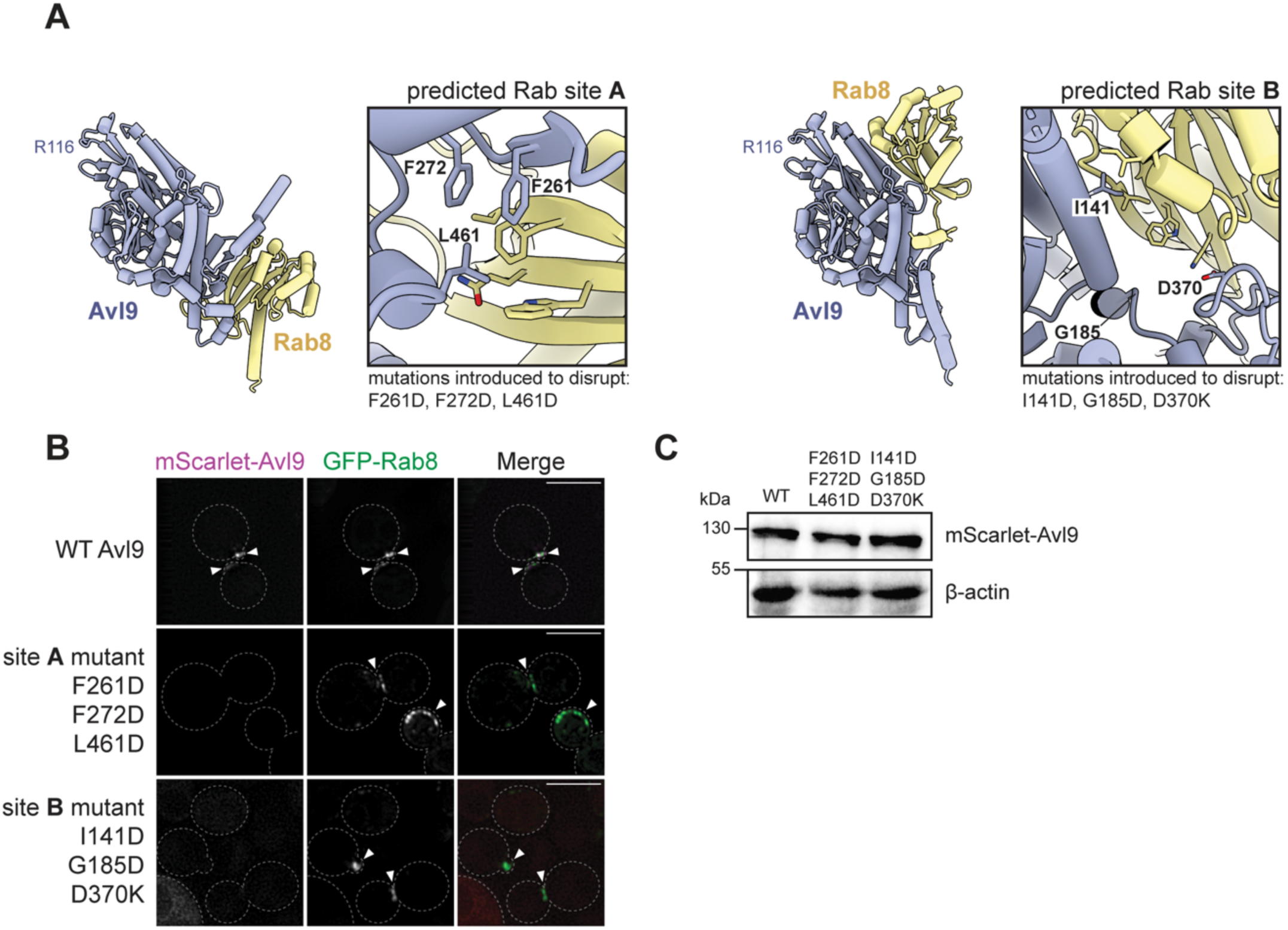
Avl9 is a Rab8 effector on secretory vesicles. (**A**) AlphaFold2 (left) and AlphaFold3 (right) structural predictions of Avl9 and Rab8. The unstructured C-terminus of Avl9 is not shown for clarity. Boxes to the right of each structure highlight the protein-protein interface with Avl9 residues labeled. (**B**) Live cell imaging of yeast expressing GFP-Rab8 and wild-type or mutant mScarlet-Avl9. Scale bars represent 5 μm. (**C**) Immunoblot of yeast whole-cell lysates.

### Human AVL9 GAP function stimulates cancer cell migration

Avl9 is well conserved throughout eukaryotes. Structural predictions of potential interactions between human AVL9 and human small GTPases are consistent with the predictions for the yeast proteins: an Arf GTPase is predicted to bind AVL9 in the equivalent surface containing a conserved arginine residue (R111) and this arginine residue is predicted in close proximity to the GTP-binding pocket of the Arf (**Figures 5A, S4A, and S4B**). Likewise, a Rab GTPase is predicted to bind AVL9 on a surface outside the catalytic domain — both AlphaFold2 and AlphaFold3 predicted that human Rab GTPases bind on the surface of human AVL9 equivalent to the predicted Rab site B of yeast Avl9 (**Figures S4A, S4B, and S4C**).

**Figure 5.**
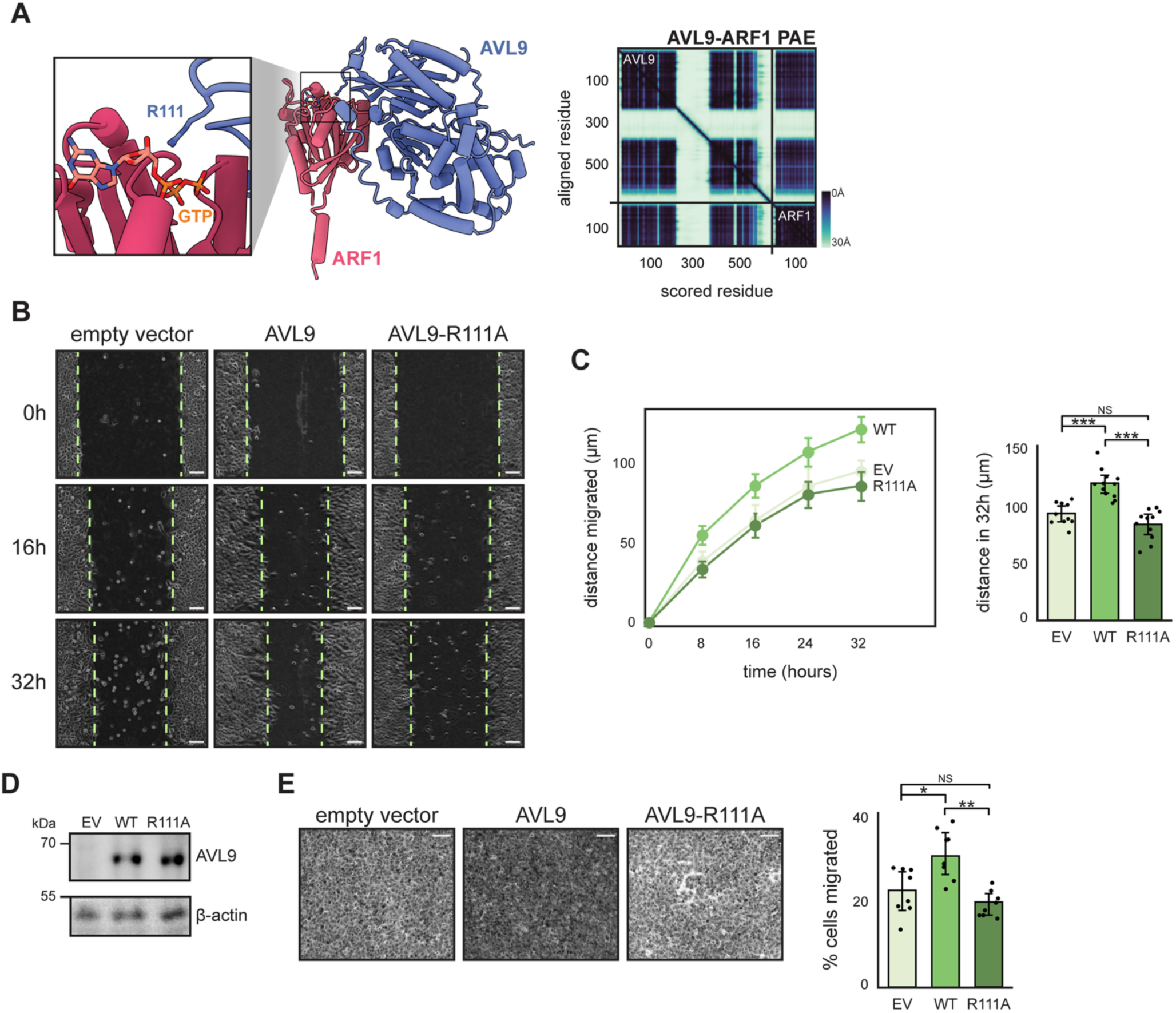
Human AVL9 GAP function stimulates cell migration. (**A**) *Left,* structural prediction of human ARF1 and human AVL9 (left), with GTP position overlaid from the crystal structure of Arf1-GTP^36^. Zoom in shows the predicted catalytic arginine (R111) of AVL9 located near GTP. *Right,* predicted alignment error (PAE) plot of AVL9-ARF1 prediction. (**B**) Representative images and quantification of wound healing assays. Dashed lines indicate wound edges and scale bar lengths represent 100 μm. Detached cells were ignored. For all experiments *AVL9* KO cells were transfected with pcDNA3 empty vector, wild-type AVL9, or AVL9-R111A. (**C**) *Left,* mean distance migrated for each experimental group over 32h period, measured as half the wound width. Data points represent mean distance migrated and error bars represent 95% confidence intervals. *Right,* quantification of total distance migrated in 32h. Data points represent measurements from replicate experiments and error bars represent 95% confidence intervals. Comparisons of total distance migrated between EV-WT (t=5.53, df=19.9, p<0.0001), R111A-WT (t=6.55, df=20.5, p<0.0001), EV-R111A (t=1.82, df=18.2, p=0.085) (Welch’s modified t-test). EV: *n*=10; WT: *n*=12; R111A: *n*=11. (**D**) Immunoblots verifying stable expression of AVL9 from pcDNA3 vectors. (**E**) *Left*, representative images of transwell migration assays. Cells that migrated through membrane pores were stained with crystal violet. Scale bars represent 100 μm. *Right*, quantification of the number of cells that migrated through pores after 24 hours. Data points represent measurements from replicate experiments and error bars represent 95% confidence intervals. Comparisons of cell migrated between EV-WT (t=2.83, df=13.9, p=0.0135), R111A-WT (t=4.75, df=10.5, p=0.0007), EV-R111A (t=1.41, df=11.0, p=0.187) (Welch’s modified t-test). Each condition *n*=8.

To determine whether GAP function is conserved in human AVL9 we made use of the observation that expression levels of AVL9 are positively correlated with cell migration^23,26^. We generated an AVL9 knockout (KO) line using CRISPR-Cas9 in A549 lung cancer cells. We transfected these *AVL9* KO cells with overexpression vectors encoding either wild-type AVL9 or AVL9 with a mutation to its putative catalytic arginine (R111A) and assessed their migration ability.

We performed scratch wound healing assays and measured the distance cells migrated into the wound. Consistent with prior studies, we found a higher amount of cell migration in cells transfected with wild-type AVL9 compared to the empty vector control (**Figures 5B and 5C**). Importantly, this effect was not observed in cells expressing the putative catalytic mutant AVL9-R111A despite similar expression levels (**Figure 5D)**.

As a complimentary approach we carried out transwell migration assays in which cells can traverse a porous membrane when exposed to a chemical gradient. We found that a greater number of cells traversed the membrane when expressing wild-type AVL9 compared to AVL9-R111A or empty vector (**Figure 5E**). Together these results support the prediction that human AVL9 functions as a GAP and indicate that the role of AVL9 in cell migration is directly tied to its GAP activity.

### Identification of human DENND6A as a GAP for ARL8B

Avl9 was not previously identified as an Arf-GAP because it does not possess the canonical Arf-GAP domain present in all known monomeric Arf-GAPs^56,57^ (**Figure S5A**), and monomeric DENN domain proteins like Avl9 have been assumed to function as Rab-GEFs^27,28,30,40^. Importantly, the predicted structure of the Avl9-Arf1 interaction is distinct from that of the known interaction between DENND1B and its GEF substrate Rab35^58^, with different surfaces of the DENN domain interacting with each GTPase (**Figure S5B**).

In contrast to the monomeric DENN domain proteins, three multimeric complexes that contain pairs of DENN domain protein subunits are known to function as GAPs: GATOR1 (a Rag-GAP)^59^, FLCN:FNIP (a Rag-GAP)^60,61^, and C9ORF72:SMCR8 (an Arf-GAP)^62,63^. Despite similar functions as GAPs, the predicted structure of the Avl9-Arf1 complex highlights key structural differences in the architectures of the monomeric and multimeric DENN domain proteins. In the multimeric complexes, two DENN domain protein subunits each contribute their longin subdomain for substrate binding and catalysis^60,62,64^ (**Figure S5C)**, which is why these complexes have been termed ‘longin domain GAPs’^65^. In contrast, while Avl9 is also predicted to use its longin domain for catalysis, its cDENN subdomain is predicted to bind to the switch regions of the GTPase substrate. Another important distinction is that the two DENN domain subunits of the ‘longin domain GAPs’ adopt very different 3D structures than the monomeric DENN domain Rab-GEF protein DENND1B (**Figure S5D**). Therefore, the predicted structure of Avl9 closely resembles the structures of the DENN domain Rab-GEF DENND1B but is distinct from the DENN domain subunits of the longin domain GAPs.

We considered the possibility that Avl9 may in fact be a subunit of a larger complex, akin to the multimeric longin GAP complexes. However, a native purification of endogenous Avl9 from *S. cerevisiae* did not yield additional co-purifying proteins (**Figure S6**), and there were no strong hits aside from GTPases in the proteome-wide predictive screen using Avl9 as a bait (including a Avl9-Avl9 homodimer). These observations are most consistent with Avl9 functioning as a monomer.

We reasoned that other monomeric DENN domain proteins may also function as GAPs. We generated pairwise structural predictions of all yeast and human DENN domain proteins with Ras-related GTPases. Predictions of yeast Anr2 and Afi1 yielded structures where conserved arginine residues of these proteins were pointed towards the nucleotide-binding pocket of an Arf GTPase (**Figures 6A and S7A**). The same was true for three human proteins, DENND11, DENND6A, and DENND6B (**Figure 6A**). In each of these structural predictions, like the Avl9-Arf1 prediction, the predicted catalytic arginine residues of these DENN proteins are located between strands β4 and β5 of their longin subdomain within a highly conserved loop, resembling an arginine finger (**Figure S7B**).

**Figure 6.**
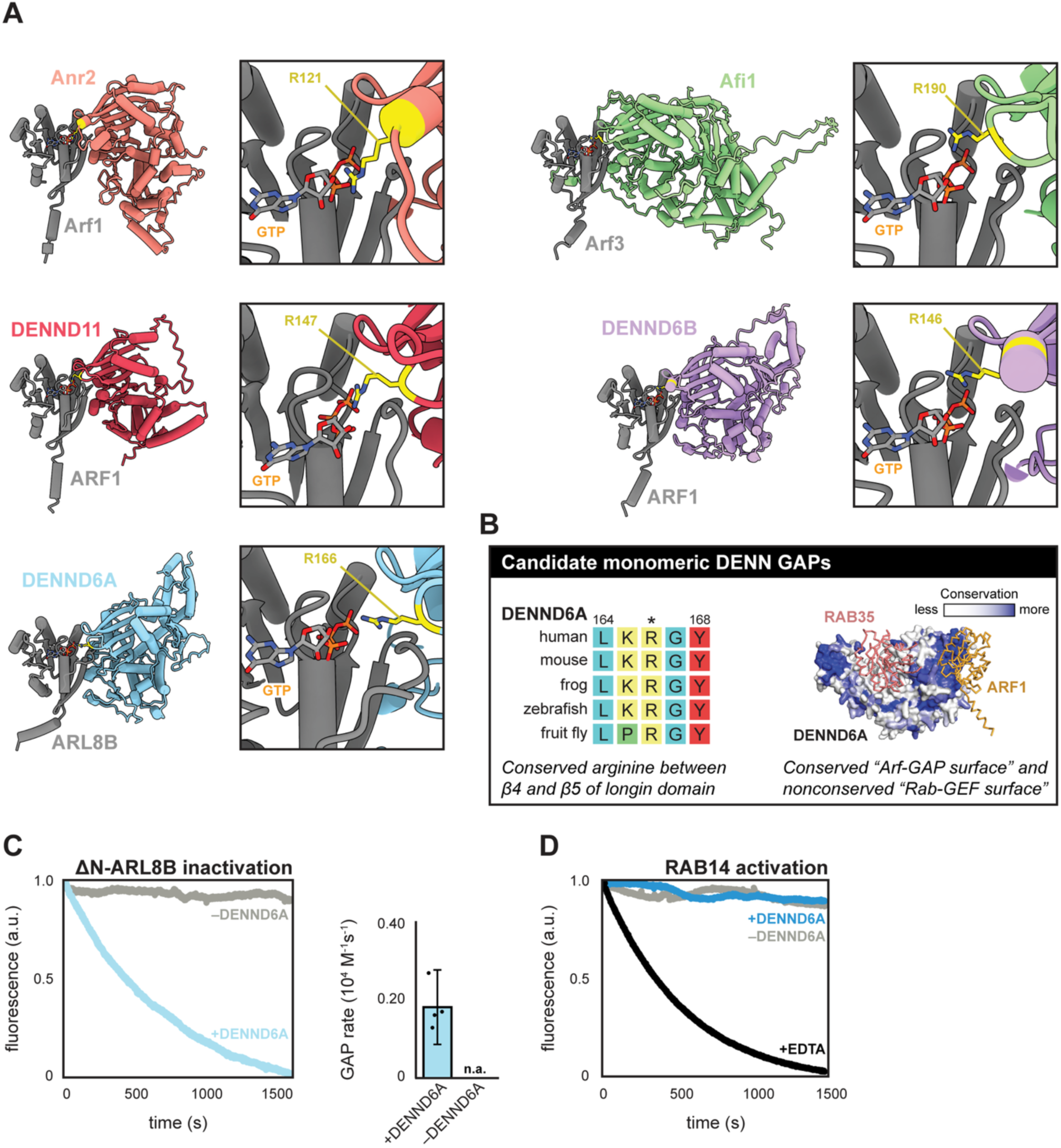
Multiple DENN domain proteins are candidate GAPs. (**A**) Structural predictions of candidate DENN GAPs interacting with GTPases. Boxes to the right of each structural prediction highlight predicted catalytic arginine residues of corresponding DENN domain protein (yellow) located near GTP, which was overlaid from a crystal structure of Arf1-GTP^36^. Arf1 is shown for each prediction except for complexes with Afi1 and DENND6A, based on results and rationale described in Discussion. Disordered termini are hidden for clarity. (**B**) Characteristics of proposed monomeric DENN GAPs. *Left,* multiple sequence alignment of DENND6A homologs. Numbers indicate residue numbers of human protein and asterisk (*) indicates putative catalytic arginine. *Right,* predicted DENND6A structure with conserved residues determined by Consurf. RAB35 and Arf1 overlaid on DENND6A structure based on alignment to known Rab-GEF (DENND1B) and Arf-GAP (Avl9). (**C**) Native tryptophan fluorescence GAP assay with ΔN-ARL8B. *Left,* representative fluorescence traces for reactions with DENND6A or mock (a.u. = arbitrary units). *Right,* quantification of GAP rate; no rate could be assigned to mock reactions (n.a.). Data points represent rates from replicate experiments and error bars represent 95% confidence intervals. (**D**) mantGDP fluorescence GEF assay with RAB14-His. Shown are representative traces for reactions with either 1 μM DENND6A, without DENND6A (mock), or EDTA (positive control).

To gain additional information regarding potential function, we next examined the evolutionary conservation of potential binding sites on each of the DENN domain proteins. As expected, we observed that the GTPase binding site used by the Rab-GEF DENND1B is highly conserved among its homologs and a subset of other DENN domain proteins, but the equivalent location is not conserved in Avl9. (**Figures 8A and 8B**). In contrast, the surface of Avl9 predicted to bind Arf1 and the loop harboring the proposed catalytic arginine residue is highly conserved (**Figure 8C**). We determined that DENND6A, DENND6B, DENND11, Afi1, and Anr2 all exhibit strong conservation at locations equivalent to the Avl9 Arf-GAP surface, and a lack of conservation at locations equivalent to the DENND1B Rab-GEF surface (**Figure 8C**). We therefore consider these to be strong candidate GAPs (**Figure 6B**).

The prediction that DENND6A is a GAP was surprising because this protein (also named FAM116A) has been reported to function as a GEF for RAB14 on recycling endosomes^26^ and for RAB34 on lysosomes^66^. DENND6A was also proposed to be an effector of the Arf-family GTPase ARL8B because it preferentially interacted with the GTP-locked mutant form of ARL8B^66^. Based on this observation, and in light of our structural predictions, we hypothesized that DENND6A instead functions as an ARL8B GAP, because GAPs also tend to interact well with the GTP-locked mutant forms of their substrates^67–70^. We therefore purified human DENND6A and a soluble variant of human ARL8B and performed *in vitro* GAP assays (**Figure S9A**). We observed that DENND6A exhibits significant GAP activity towards ARL8B, with a rate of 0.18 x 10^4^ M^-1^s^-1^ (**Figure 6C**), which is comparable to the GAP rate of Age2 towards Arf1 (**Figure 2F**).

Importantly, reported cellular phenotypes provide strong support for DENND6A functioning as an ARL8B GAP in cells. ARL8B is known to play a key role in lysosome positioning by linking lysosomes to kinesin-1^71^. Accordingly, ARL8B knockdown led to accumulation of lysosomes in the perinuclear region while overexpression of ARL8B resulted in movement of lysosomes to the periphery. In contrast, DENND6A perturbation led to the opposite phenotypes: knockdown of DENND6A resulted in lysosome dispersion to the cell periphery while overexpression resulted in lysosome clustering near the nucleus^66^. These are the expected results for a negative regulator of ARL8B function in lysosome positioning and provide a physiological function for DENND6A GAP activity in cells.

The published biochemical evidence in support of DENND6A Rab-GEF function involved assays in which nucleotide exchange activity appeared to be low and kinetic rates were not determined^26,66^. We therefore purified human RAB14 for use in GEF assays which utilize a fluorescent analog of GDP to monitor nucleotide exchange in real time^72^. Despite multiple attempts under different conditions, we could not detect GEF activity towards RAB14 (**Figures 6D and S9B**). We observed nucleotide exchange at high RAB14 concentrations, but this was independent of DENND6A and therefore represents intrinsic exchange. Although it is possible that different conditions are needed for activity than the ones we employed, the absence of GEF activity is consistent with the lack of conservation of the expected Rab-GEF binding site in DENND6A (**Figure S8C**). Taken together, the available results provide strong support for a role of DENND6A as a GAP for the Arf-family GTPase ARL8B both *in vitro* and in cells.

## DISCUSSION

We identified Avl9 in a computational screen for interactors of Arf1. This interaction was not identified in previous large-scale screens^18–21,76–79^, highlighting the utility of predictive screens focusing on specific target proteins of interest. Avl9 was first characterized as a factor involved in secretory transport from the Golgi^22^. Yeast with deletions of *VPS1* (dynamin-like GTPase localized at endosomes) and *APL2* (subunit of the AP-1 complex) exhibit synthetic lethality with deletion of *AVL9*. Avl9 is conserved throughout eukaryotes and dysregulation of human AVL9 has been implicated in several cancers^23–25^. Knockdown of AVL9 results in a decrease in cell migration and, conversely, overexpression leads to an increase in cell migration^26^. Despite its involvement in the secretory pathway and relevance to human disease, the molecular function and biological role of Avl9 was unknown.

We performed a series of genetic and biochemical experiments to determine the physiological substrate(s) of Avl9. Avl9 exhibited highest GAP activity towards Arf1 but acted on each of the Arf-family GTPases tested. Promiscuity appears common in GAPs^41,42^, and it is important to note that Avl9 did not possess Rab-GAP activity. Given its role in trafficking at the trans-Golgi network, Arf1 is the most likely substrate *in vivo*, but it is also possible that Avl9 functions to ensure GTPase quality control by ‘erasing’ any Arf GTPases that may leak onto Golgi-derived secretory vesicles.

We also determined that Avl9 is recruited to secretory vesicles through an interaction with Rab8, representing GTPase crosstalk (**Figure 7**). Rab8 appears capable of binding to two sites on Avl9, but site B appears to be more important and conserved. Interestingly, Harsay *et al.* identified a deleterious *AVL9* mutation (*avl9*-G52D)^22^ that is predicted to disrupt site B. It is possible that human AVL9 localizes via interaction with a different Rab GTPase, as it was found to localize to recycling endosomes, which are primarily occupied by Rab11^26,80^. Recently, Stockhammer *et al*. characterized Arf1 compartments that mature into recycling endosomes, a process defined by the shift from Arf1 to Rab11 occupancy^81^. The authors predicted that Rab11 recruits an Arf-GAP to facilitate the Arf1 to Rab11 transition and we speculate that AVL9 may fulfill this role.

**Figure 7.**
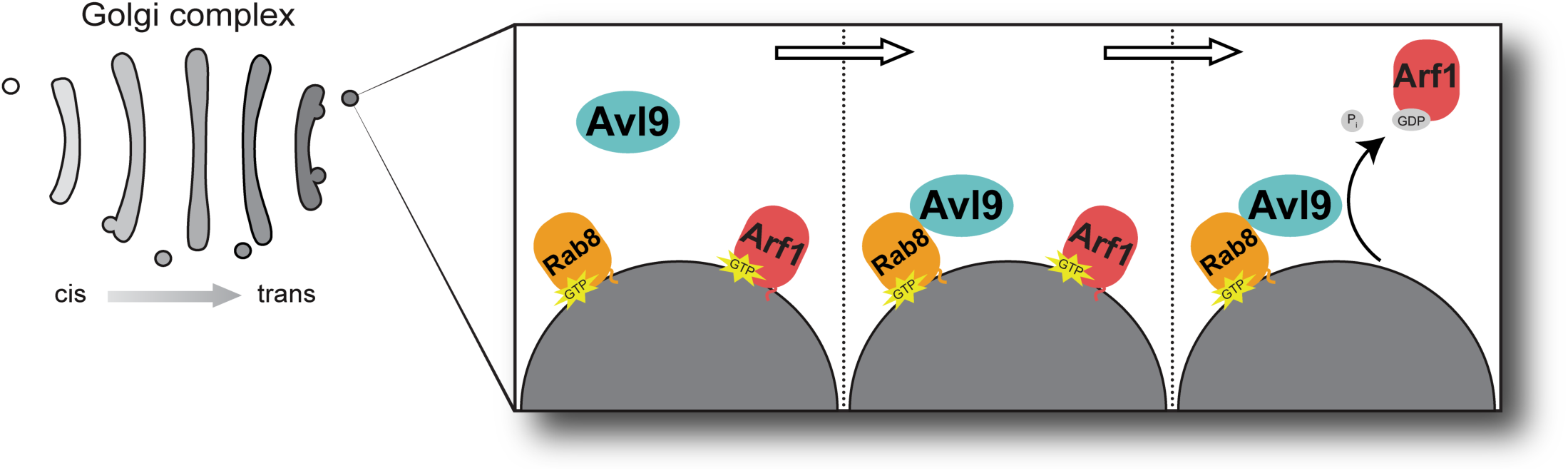
Model of Avl9 function in secretion. Arf1 sets in motion the steps necessary for its own release from secretory vesicles via GTPase crosstalk. Arf1 recruits the Rab-GEF TRAPPII which activates Rab11 (encoded by yeast *YPT31* and *YPT32*)^73^. Rab11 in turn recruits the Rab-GEF Sec2 which activates Rab8 (encoded by yeast *SEC4*)^74,75^. As shown, Rab8 then recruits Avl9 to secretory vesicles where it inactivates Arf1, ‘erasing’ it from the vesicle.

Human AVL9 was previously shown to play a role in cultured cancer cell migration^23,25,26^ and we have recapitulated these findings using both wound healing and transwell migration assays. We found that mutation of the predicted catalytic arginine of AVL9 phenocopied empty vector controls, suggesting that GAP activity is tied to cell migration via regulation of Arf activity^82^. As vesicular trafficking and cell migration are highly intertwined processes^83,84^, it is likely that both the enzymatic function and biological role of AVL9 are conserved. Future studies are needed to determine precisely how AVL9 modulates cell migration.

Our discovery that Avl9 is an Arf-GAP raised the possibility that other monomeric DENN domain proteins may function as GAPs. Based on our analyses we propose that monomeric DENN domain proteins with a conserved surface on their longin subdomain that includes an arginine finger between strands β4 and β5 are DENN GAPs. In addition to yeast and human AVL9 we identified several candidate DENN GAPs: DENND6A, DENND6B, and DENND11 in humans, and Afi1 and Anr2 in yeast. We suspect that yeast Arf3 may be the substrate of Afi1, based on a previously published study in which Arf3 localization was perturbed with loss of Afi1 and Afi1 interacted preferentially with a GTP-locked variant of Arf3^85^. Similarly, DENND6A was previously shown to preferentially interact with the GTP-locked mutant form of ARL8B^66^. We selected DENND6A for further investigation and determined that it exhibits strong GAP activity towards ARL8B. This finding provides a molecular mechanism for the lysosomal distribution phenotypes of cells lacking or overexpressing DENND6A^66^.

DENND6A was previously reported to act as a Rab-GEF, and its loss resulted in altered localization patterns of RAB14 and RAB34 in cells. However, in each case the Rab GTPase maintained significant localization to recycling endosomes and lysosomes, respectively, which would not be expected if DENND6A was the GEF for these Rabs at these locations^26,66^. The observed mislocalization of these Rabs can be explained instead by disrupted organellar trafficking and transport caused by hyperactivation of ARL8B, and potentially other closely related GTPases, upon loss of DENND6A GAP function.

The localizations of the other candidate monomeric DENN GAPs have been reported: DENND6B shares strong sequence similarity with DENND6A and localizes to recycling endosomes^26^; Afi1 localizes to the plasma membrane^85^; Anr2 localizes to lipid droplets^86^; and DENND11 localizes to the Golgi^87^. This study now enables future work aimed at identifying the GTPase substrates of these proteins and dissecting how they regulate these organelles.

As we have determined that two monomeric DENN domain proteins in yeast and humans possess robust GAP activity towards Arf-family GTPases, the functional repertoire of this protein family is greater than previously appreciated. Our results indicate that monomeric DENN domain proteins should not be assumed by default to function as Rab-GEFs, as several belong instead to a distinct class of monomeric ‘DENN GAPs’ defined by Avl9. Candidate members of this family are identifiable by the presence of a conserved patch containing an arginine finger on the surface of the longin subdomain (**Figure 6B**).

## METHODS

### Yeast strains and plasmids

All strains (**Table S1**) and plasmids (**Table S2**) were constructed using conventional methods and are available upon request. Yeast shuffling assays were performed by transforming CFY5184 (already containing pRS416-APL2) with either empty pRS415 vector or pRS415 vector encoding Avl9 variants. Cells were grown overnight in synthetic complete medium (SC) lacking tryptophan, serially diluted in 96 well plates, and pinned to SC agar lacking tryptophan (control) and SC agar containing 1 mg/mL 5-fluoroorotic acid (5-FOA).

### *In silico* protein-protein interaction screens

Protein-protein interaction predictions were performed using custom scripts to run the LocalColabFold (github.com/YoshitakaMo/localcolabfold) implementation of AlphaFold2 (ColabFold v1.5.5 and AlphaFold v2.3.2)^16,17,88^. Primary sequence of a ‘bait’ protein was combined pairwise with the primary sequence of ‘prey’ proteins in a .csv file format readable by ColabFold. The list of prey proteins was comprised of approximately 450 Golgi and post-Golgi secretory pathway proteins (determined by GO term annotations and *Saccharomyces* Genome Database gene descriptions^43^) or the entire yeast proteome. Three predictions were generated for each protein-protein input, with templates, 5 recycles, and a recycle early-stop tolerance of 0.5. Predictions were ranked by averaged ipTM scores. We considered a predicted interaction plausible when ipTM≥ 0.6; plausible predictions were manually inspected by viewing PAE plots and PDB files (PyMol v2.5.5 or ChimeraX v1.9). Scripts for this pipeline are accessible at github.com/FrommeLab/colabfold-pipeline.

### Protein purification

*S. cerevisiae* Avl9 and human DENND6A were purified from *Pichia pastoris* expressing GST fusion proteins under control of the *AOX1* promoter. Cells were cultured in 5 mL BMGY media (0.8% yeast extract, 1.6% peptone, 0.24% K_2_HPO_4_, 1.17% KH_2_PO_4_, 1.34% yeast nitrogen base, 0.4 μg/mL biotin, 1% glycerol) at 30°C overnight. Cultures were diluted 1:40 into 200 mL fresh BMGY. After 8 hours, cultures were diluted 1:40 into 1 L autoinduction media (identical to BMGY, but with 0.5% methanol and 0.4% glycerol) and grown at 25°C. Additional methanol (to a final concentration of 0.5%) was added to the cultures twice: 16 hours- and 40 hours-post dilution into autoinduction media. At 64 hours post-dilution, cells were collected by centrifugation, pellets were resuspended 1:1 in lysis buffer (50 mM Tris pH 7.6, 300 mM NaCl, 10% glycerol, 0.5% CHAPS, 1 mM DTT, plus a protease inhibitor cocktail (Roche 11836145001)), frozen dropwise in liquid nitrogen, and lysed in a SPEX cryogenic mill. Lysates were thawed on ice and clarified by centrifugation at 47,000 x g. Clarified lysates were then incubated with glutathione resin (Cytiva 17075601) for 2 hours at 4°C with gentle agitation. Resin was washed three times with lysis buffer and once with DENN storage buffer (50 mM Tris pH 7.6, 150 mM NaCl, 10% glycerol, 0.1% CHAPS, 1 mM EDTA, 1 mM DTT). Protein was eluted by incubating with PreScission protease for ∼16 hours at 4°C.

Myristoylated yeast Arf1, ΔN-Arf GTPases (Arf1, Arf3/ARF6, Arl1, Arl3/ARFRP1, Sar1), His-Rab GTPases (Sec4/Rab8, Vps21/Rab5, Ypt31/Rab11), prenylation proteins (Bet2, Bet4, Mrs6/REP), Gdi1/GDI, and full-length Sec4/Rab8 were purified as previously described^73,89,90^. Human ΔN-ARL8B and human RAB14-His were purified in the same manner as yeast Arf and Rab GTPases. Briefly, Rosetta2 *E. coli* (Novagen 71397) were transformed with expression vectors and grown in terrific broth at 37°C to an OD of ∼3.0. Temperature was reduced to 18°C and expression was induced overnight with 300 μM IPTG. Cells were lysed by sonication in 5 mL lysis buffer (30 mM Tris pH 7.6, 300 mM NaCl, 5% glycerol, 2 mM MgCl_2_, 1 mM DTT, plus a protease inhibitor cocktail) per 1 g of cell pellet. Lysate was clarified by centrifugation at 47,000 x g. Proteins were affinity-purified using glutathione resin (Rabs) or Ni-NTA resin (Arfs) (Thermo Scientific 88223). Rabs were eluted by incubating with PreScission protease for ∼16 hours at 4°C. Arfs were eluted using lysis buffer plus 500 mM imidazole, His-tag was cleaved via overnight incubation with TEV protease at room temperature, and then buffer exchanged into lysis buffer without imidazole using Zeba spin desalting columns (Thermo Scientific 89877).

Endogenous Avl9 was purified from a TAP-tag *S. cerevisiae* strain (Horizon Discovery YSC1178-202232335) using previously-described tandem affinity methods^91,92^. Briefly, 12L of yeast were grown at 30° in YPD to an OD of ∼2. Cells were pelleted, resuspended 1:1 in lysis buffer (50 mM Tris pH 7.6, 300 mM NaCl, 10% glycerol, 0.5% CHAPS, 1 mM DTT) plus a protease inhibitor cocktail tablet, frozen dropwise in liquid nitrogen, and lysed in a SPEX cryogenic mill. Lysate was clarified via centrifugation at 47,000 x g, affinity-purified with IgG affinity resin (Cytiva 17096901), and eluted by incubation with TEV protease. The elution was further purified by binding to calmodulin affinity resin (Agilent 214303) and eluted with calmodulin elution buffer (25 mM Tris pH 8.0, 300 mM NaCl, 5% glycerol, 0.1% CHAPS, 1mM magnesium acetate, 1mM imidazole, 20 mM EGTA, 1 mM DTT).

All purified proteins were aliquoted, snap frozen in liquid nitrogen, and stored at -80°C.

### Prenylated Rab8-GDI complex preparation

Prenylated Rab8-GDI complex was prepared as previously described for other yeast Rabs^73,92^. Briefly, 40 μM full-length Sec4/Rab8 was incubated with 200 μM GDP and 20 mM EDTA in prenylation buffer (20 mM HEPES, 150 mM NaCl, 2 mM MgCl_2_, 1 mM DTT) for 30 min at 30°C. MgCl_2_ was added to a final concentration of 25 mM to halt the exchange reaction. Excess EDTA and MgCl_2_ were removed by buffer exchanging with Zeba spin desalting columns. Rab8 prenylation was achieved by combining Sec4/Rab8, Gdi1, 6xHis-Bet2-Bet4, and 6xHis-Mrs6 in a 10:10:1:1 ratio in prenylation buffer, with 120 μM geranyl-geranyl pyrophosphate (Millipore Sigma G6025) and 25 μM GDP. After 1 hour at 37°C, Ni-NTA resin was added to remove geranylgeranylation proteins and Rab8-GDI complex was isolated by gel filtration chromatography.

### Liposome preparations

Synthetic ‘Golgi’ liposomes were prepared with a composition of lipids mimicking that of the late-Golgi/TGN-derived vesicles^93^. Lipids solubilized in chloroform were mixed in the molar ratios described in **Table S3**. The lipid mix was vacuum dried and rehydrated in HK buffer (20 mM HEPES pH 7.4, 125 mM KOAc) overnight at 37°C. Lipids were extruded through either 100 nm filters (GAP and GEF assays) or 400 nm filters (pelleting assays) and stored at 4°C. Golgi Ni^2+^ liposomes also included 5% Ni^2+^-DOGS.

### Liposome pelleting assays

To assess Rab-mediated membrane recruitment of Avl9, liposome pelleting was performed as described previously^73^. 7.5 μM His-Rab was activated on 600 μM Golgi Ni^2+^ liposomes in HK buffer with 125 μM GTP and 1 mM EDTA, in a polyallomer tube, for 30 min at room temperature. Nucleotide exchange was halted by the addition of 2 mM MgCl_2_. Avl9 was added to a final concentration of 1 μM and the tube was incubated for 15 min at room temperature. Samples were centrifuged at 150,000 x g for 15 min at 4°C and, subsequently, the liposome pellet was separated from supernatant. Pelleting assays with prenylated Rab8 was performed identically but with Golgi liposomes and 5 μM prenylated Rab8-GDI complex. Samples were analyzed via SDS-PAGE; proteins were visualized by Coomassie and lipids were analyzed by DiR dye. Analyses were performed in ImageJ (v1.8.0_172). 4-5 technical replicates were performed for each condition.

### Malachite green GAP assays

GAP activity of Avl9 towards each of the yeast Arf GTPases was initially measured using endpoint colorimetric assays. GTPases were activated by incubation with fivefold molar excess of GTP and twofold molar excess EDTA over MgCl_2_ for 1 hour at room temperature. Exchange was halted by the addition of excess MgCl_2_. GTP-loaded GTPases were buffer exchanged into HKM buffer (20 mM HEPES pH 7.4, 125 mM KOAc, 1 mM MgCl_2_) using Zeba spin desalting columns. 12 μM GTP-loaded GTPase was added to a PCR tube followed by 3 μM Avl9 and HKM buffer to a final volume of 40 μL, all on ice. Reactions then proceeded for 5 min at 30°C. Reactions were transferred to wells of a 96-well plate containing 200 μL of malachite green reagent (Millipore Sigma MAK113A). Plates were left at room temperature for 30 min before measuring absorbance at 620 nm with a BioTek Synergy H1 microplate reader. Mock reactions were performed by adding Avl9 storage buffer in place of Avl9. Concentration of free phosphate produced was determined by a standard curve generated from phosphate standards (Millipore Sigma MAK113B). Avl9 activity towards each GTPase was calculated by the following equation: enzyme activity = (P_Avl9_ − P_mock_) / time, where P is the amount of free phosphate (in μM) produced from reactions with Avl9 (P_Avl9_) or without (P_mock_). At least 8 technical replicates were performed for each Arf GTPase and 6 technical replicates were performed for Rab5.

### Tryptophan fluorescence GAP assays

The kinetic rate of GAP activity of Avl9 towards Arf1 was measured using previously-described methods, where the nucleotide-bound state of Arf1 was monitored via native tryptophan fluorescence (297.5 nm excitation, 340 nm emission) using a PTI fluorometer^44,45^. 4.0 μM myristoylated Arf1 and 0.75 μM prenylated Rab8 (in complex with GDI until activated) were first activated on 400 μM Golgi liposomes in HKM buffer by adding 100 μM GTP and 2 mM EDTA, followed by a 10 min incubation. To quench exchange, 5 mM MgCl_2_ was added. Avl9 was then added to a final concentration of 1 nM and measurements began immediately. For reactions where Arf1, Rab8, or Avl9 was excluded, protein storage buffer was added in its place. All steps were performed at 30°C. Fluorescence traces were fit to single-exponential decay curves to determine the rate constant (k_GAP_). GTP hydrolysis rates were then calculated by the equation: rate = k_GAP_ / [GAP]. The traces shown in figures have been normalized using the fluorescence span of the reaction. Analyses were performed in R (v4.2.1). GAP assays with ΔN-Arf1 or ΔN-ARL8B were performed similarly, except that no liposomes nor Rab were included and 1 μM GAP (Avl9, Age2, or DENND6A) was used.

### mantGDP fluorescence GEF assays

RAB14-7xHis was loaded with mantGDP (Invitrogen M12414) by incubation with fivefold molar excess of mantGDP and twofold molar excess EDTA over MgCl_2_ for 1 hour at room temperature. Exchange was halted by the addition of excess MgCl_2_. mantGDP-loaded RAB14 was then buffer exchanged into HKM buffer using Zeba spin desalting columns. GEF activity of DENND6A was measured using previously-described methods, where the nucleotide-bound state of RAB14 was monitored via mantGDP fluorescence (365 nm excitation, 440 nm emission)^94,95^. mantGDP-RAB14 (1.0 or 4.0 μM) and 200 μM GTP were incubated with or without 333 μM Ni^2+^ Golgi liposomes in HKM buffer. DENND6A was then added to a final concentration of 1.0 or 2.0 μM and measurements began immediately. Positive control nucleotide exchange reactions were performed by adding excess EDTA in place of DENND6A. The intrinsic exchange rate of RAB14 was measured by adding DENN storage buffer in place of DENND6A. All steps were performed at 30°C. The traces shown in figures have been normalized using the fluorescence span of the reaction.

### Fluorescence microscopy and image analysis

Yeast cells were grown at 30°C in synthetic media to mid log phase. Images were collected with a DeltaVision RT widefield deconvolution microscope. Image acquisition and deconvolution were performed in SoftWoRx software (v7.0.0). Deconvolved images were preprocessed in ImageJ (v1.8.0_172) using the Stack Box plugin (github.com/ryanfeathers/Stack_Box) to generate cropped image stacks of one focal plane containing 1-2 budding cells^92^. Colocalization analyses were performed in CellProfiler (v4.2.8).

### Cell culture

A549 lung cancer cells were cultured in F-12K medium supplemented with 10% fetal bovine serum (FBS). HEK293T cells were cultured in DMEM medium with 4.5 g/L glucose, L-glutamine, sodium pyruvate, and 10% FBS. Cells were incubated in a humidified environment at 37°C with 5% CO_2_. Lipofectamine 2000 (Invitrogen 11668027) was used for plasmid transfections following manufacturer’s instructions.

### Lentivirus production and transduction to generate AVL9 KO cells

Lentivirus was produced in HEK293T cells by transfecting 70% confluent cells with packaging plasmids, VSVg and Pax2, and lentiCRISPRv2 plasmids with gRNA sequences targeting either *AVL9* or non-targeting controls at a 1:2:3 ratio. Virus-containing medium was collected 24, 32, and 48 hours post-transfection. Collected medium was combined, passed through a 0.45 μM filter, and stored at 4°C (short term) or -80°C.

A549 cells were seeded in 6-well plates until 70% confluent. Transduction was carried out using virus medium and fresh medium at a 3:1 ratio plus 8 μg/mL polybrene (Millipore Sigma TR-1003). A “kill control” well received fresh medium alone. This transduction process was repeated every 12 hours for a total of three times, after which fresh medium was added, and cells were left for 12 hours before drug selection. Cells were trypsinized, seeded into wells of a 6-well plate, and incubated with 1.5 μg/mL puromycin until all kill control cells died. Medium was changed every two days during drug selection. Gene disruption confirmed by sequencing.

### Wound healing assays

48 hours after pcDNA3 transfection, in 24-well plates, wounds were scratched in cell monolayer using a 20 μL pipette tip. Wells were then washed twice with PBS and F-12K medium with 2% FBS was added. Plates were then incubated for 30 min at 37°C and 5% CO_2_. Timing started after this incubation period. Imaging was performed using a Nikon Eclipse Ti inverted microscope with a 10x objective lens and an Andor Neo sCMOS camera. Plates were maintained at 37°C and 5% CO_2_ throughout imaging using a Tokai Hit STX onstage incubator. Images were collected every 8 hours. Analyses were performed in ImageJ (v1.8.0_172) using the Wound Healing Size Tool (github.com/AlejandraArnedo/Wound-healing-size-tool)^96^. Images of wells where detached cells interfered with calling of wound edges were rejected.

### Transwell migration assays

48 hours after pcDNA3 transfection, cells were detached and washed twice with PBS. Cells were resuspended in F-12K medium (without FBS), live cells were counted with a hemocytometer using trypan blue, and diluted to a concentration of 1x10^6^ live cells/mL. 100 μL of the cell solution was pipetted onto an 8.0 μm pore membrane of a transwell insert (Corning 3422) placed in a 24-well plate. The plate was incubated for 10 min at 37°C and 5% CO_2_ then 600 μL of F-12K medium supplemented with 10% FBS was added to the well below the transwell insert. Plates were then incubated for 24 hours at 37°C and 5% CO_2_. Representative images were obtained after staining, using a Nikon Eclipse Ti inverted microscope and an Andor Neo sCMOS camera.

Quantification of migrated cells was performed with crystal violet staining. Medium below the transwell inserts was aspirated and non-migrated cells were removed from the upper side of the membrane with cotton swabs. 750 μL of PBS was added into the well below the insert to wash migrated cells. PBS was removed and 750 μL of ice-cold methanol was added to the well to fix cells. Following 20 min at room temperature, methanol was removed and membranes were left to air dry for 30 min. Cells were stained with 750 μL of 0.5% crystal violet by adding it to the wells and incubating for 20 min at room temperature. Transwell inserts were then washed with H_2_O until excess stain was completely removed. Transwell inserts were placed in wells containing 500 μL of 33% acetic acid for 10 min to lyse cells and release crystal violet. Transwell inserts were removed and absorbance of the solution was measured at 595 nm using a BioTek Synergy H1 microplate reader. The number of migrated cells was determined using a standard curve generated by seeding known amounts of cells in 24-well plates and following the above staining protocol.

### Protein isolation for immunoblots

Yeast cells were grown to mid-log phase and five OD_600_ worth of cells were collected, centrifuged, and washed twice with H_2_O. Pellets were resuspended in 1 mL cold H_2_O, 110 μL trichloroacetic acid was added, and samples were left on ice for 30 min. Samples were pelleted by centrifugation at 16,000 x g for 3 min at 4°C. The supernatant was aspirated, 1 mL cold acetone was added to pellets, and pellets were resuspended in a bath sonicator. Samples were pelleted and dried in a vacuum concentrator for 1 min. Pellets were resuspended in 50 μL boiling buffer (50 mM Tris pH 7.5, 1 mM EDTA, 1% SDS), vortexed for 5 min with glass beads, and heated for 5 min at 55°C. 50 μL of urea sample buffer (150 mM Tris pH 6.8, 6M urea, 6% SDS, bromophenol blue, 10% β-mercaptoethanol) was added and vortexed for 5 min. Samples were heated for 10 min at 55°C. Beads were pelleted and supernatant was collected for analysis.

A549 cells were washed in ice-cold PBS, resuspended in ice-cold RIPA buffer (50 mM Tris pH 8.0, 150 mM NaCl, 1% Igepal CA-630, 0.5% sodium deoxycholate, 0.1% SDS, 1 mM PMSF), and agitated for 20 min at 4°C. Samples were then centrifuged at 16,000 x g for 20 min at 4°C. Supernatant was collected and protein concentration of each sample was determined using a BCA assay (Thermo Scientific 23225) and normalized with RIPA buffer.

### Immunoblots

Samples were analyzed via SDS-PAGE and western blotting. Blotting of β-actin was used as a loading control. Rabbit polyclonal anti-AVL9 antibody (GeneTex GTX16209) was used at a 1:500 dilution. Rabbit polyclonal anti-RFP antibody (Rockland 600-401-379) was used at a 1:1000 dilution. Rabbit polyclonal anti-β-actin antibody (Invitrogen PA1-183) was used at a 1:5000 dilution. HRP-conjugated anti-rabbit antibody (Cytiva NXA931V) was used at a 1:10000 dilution.

## ACKNOWLEDGEMENTS

We thank J. Liu and Q. Huang for feedback on the manuscript. We thank the laboratories of J. Baskin, T. Bretscher, R. Collins, S. Emr, R. Schekman, and C. Ungermann for sharing strains, equipment, and reagents. We thank S. Huang for advice with the cell culture experiments, B. Bishop for assistance with preliminary experiments, and K. Manzer for providing Age2 protein. Wound healing imaging data were acquired through the Cornell Institute of Biotechnology BRC Imaging Facility. This work was supported by National Institutes of Health fellowship F32GM155980 to RCV and grant R35GM136258 to JCF.

## AUTHOR CONTRIBUTIONS

RCV: conceptualization, formal analysis, investigation, manuscript writing, visualization, funding acquisition JCF: conceptualization, manuscript writing, visualization, supervision, funding acquisition

---

**Figure S1.**
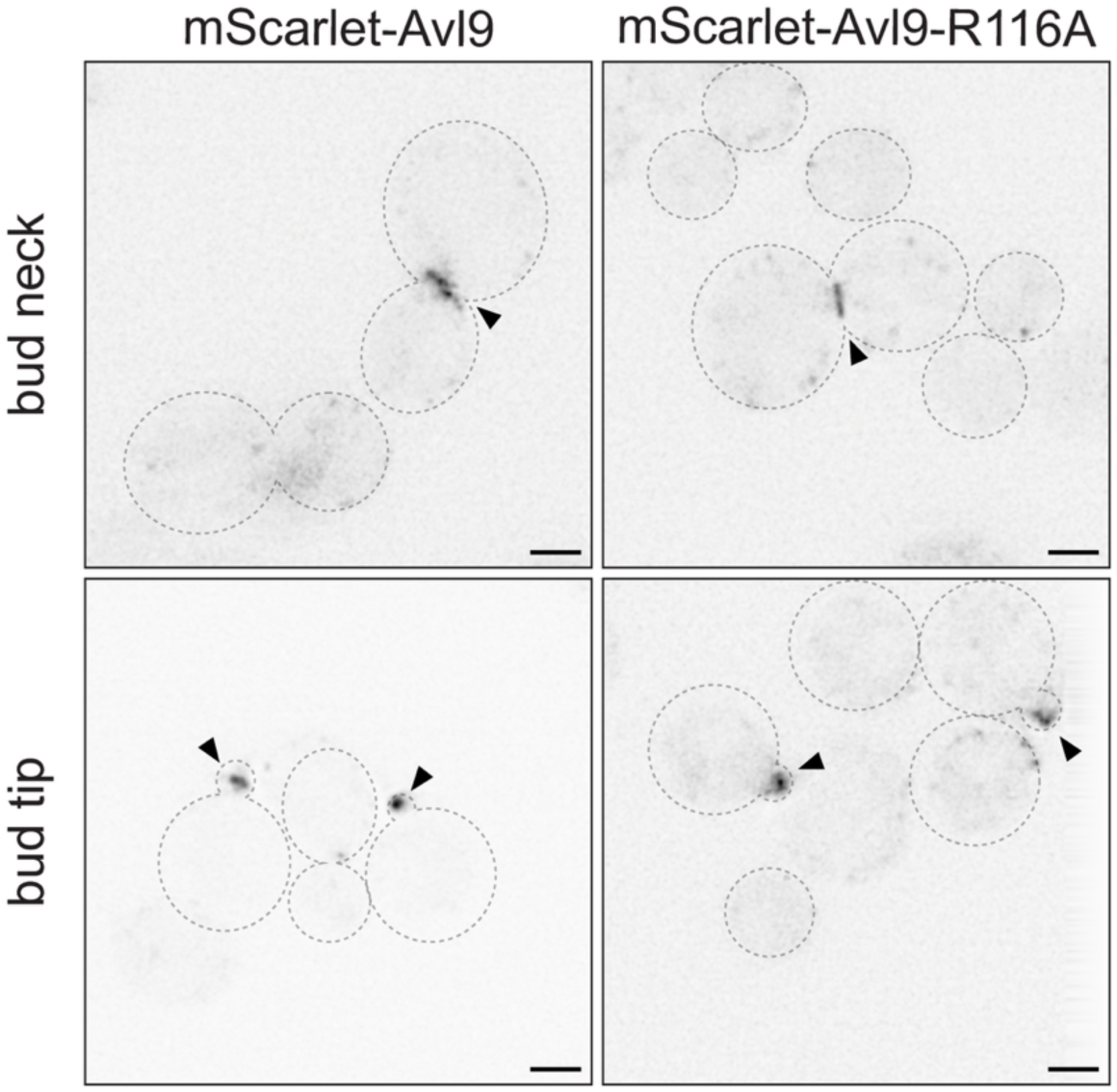
Avl9-R116A expresses and localizes similar to wild-type Avl9. Live cell fluorescence microscopy images. mScarlet-tagged Avl9 or Avl9-R116A expressed as sole copy of Avl9 from plasmids. Scale bars represent 2 μm.

**Figure S2.**
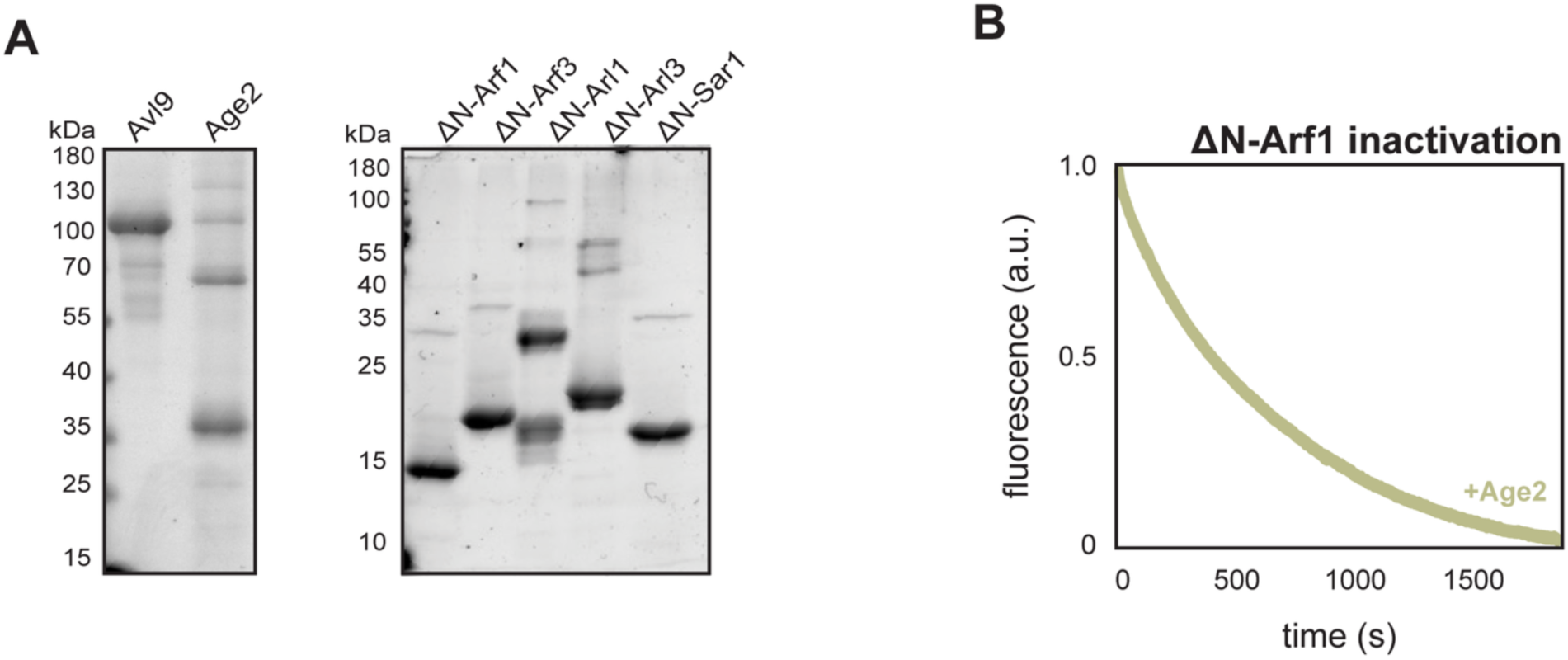
*In vitro* GAP assays. (**A**) SDS-PAGE gels stained with Coomassie showing ΔN-Arf GTPases, Avl9, and Age2 used in biochemical assays. Note that ΔN-Arl1 appears to form SDS-resistant dimers (∼35 kDa) that arise even after isolation of monomeric ΔN-Arl1 fraction via size exclusion chromatography. (**B**) Full reaction span of native tryptophan fluorescence Age2 GAP assay with ΔN-Arf1 (a.u. = arbitrary units).

**Figure S3.**
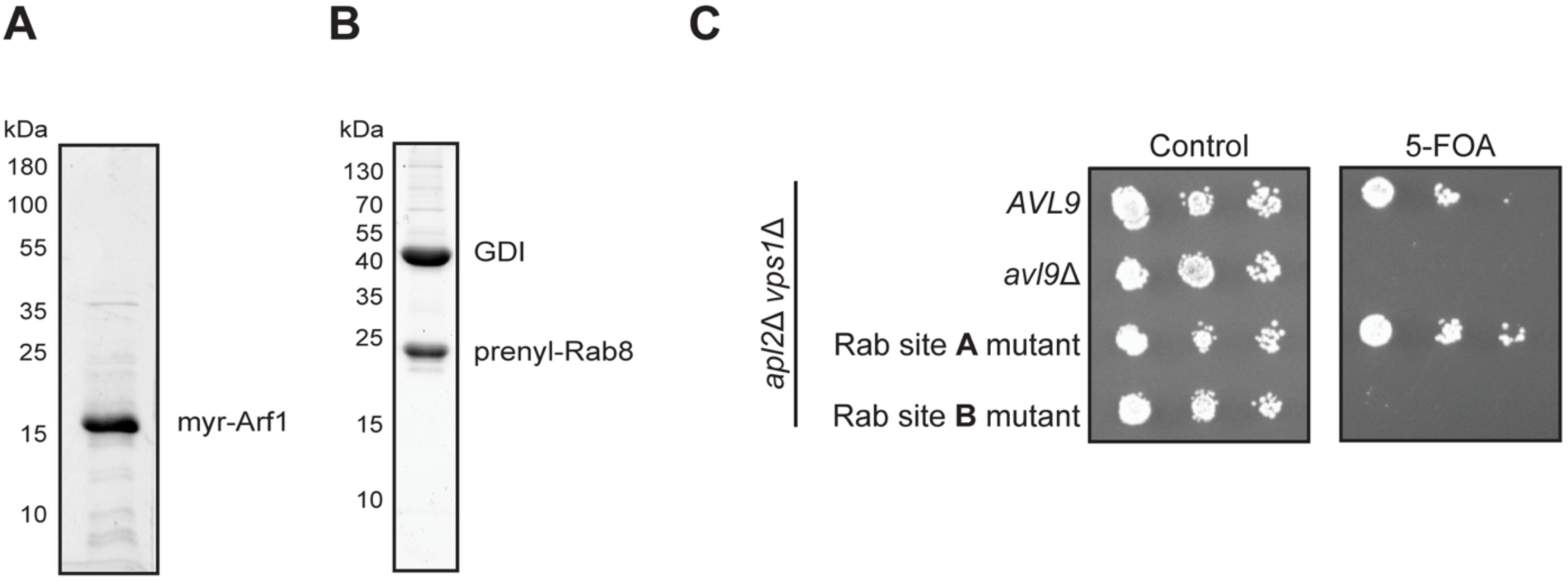
Avl9 is a Rab8 effector. (**A, B**) SDS-PAGE gels stained with Coomassie showing purified full-length, myristoylated Arf1 and prenylated Rab8-GDI complex. (**C**) Yeast growth assay, cultures were serially diluted from left to right. *URA3* plasmid encoding *APL2* counter-selected on 5-FOA. *AVL9* Rab site A mutant = F261D F272D L461D and Rab site B mutant = I141D G185D D370K.

**Figure S4.**
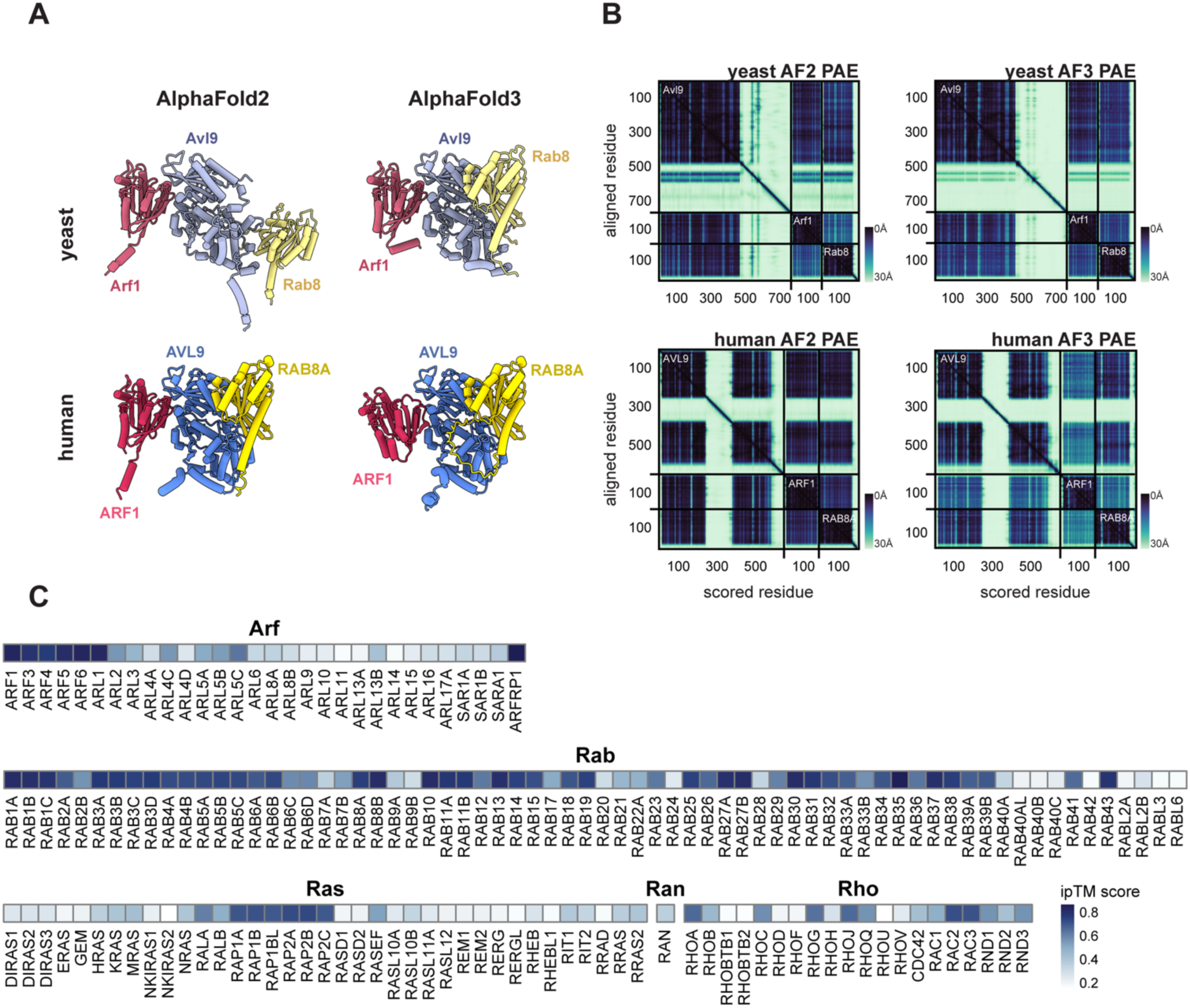
Comparison of yeast and human Avl9-Arf1-Rab8 structural predictions. (**A**) AlphaFold2 (AF2) and AlphaFold3 (AF3) structural predictions of yeast or human Avl9-Arf1-Rab8 in complex and (**B**) corresponding PAE plots. (**C**) Heatmap showing the average ipTM score of structural predictions for AVL9 and a given GTPase.

**Figure S5.**
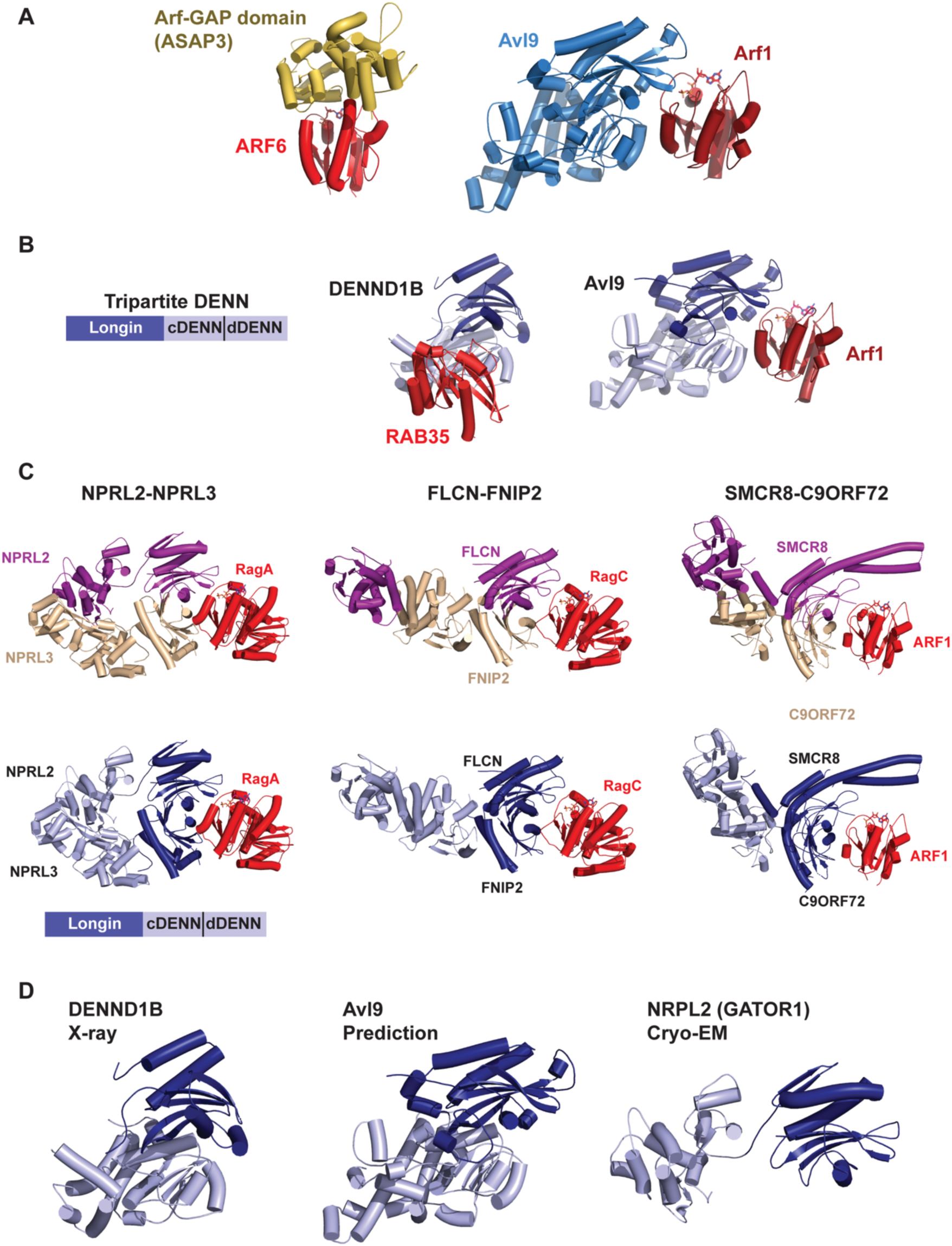
Comparison of DENN domain protein structures. (**A**) ASAP3-ARF6 X-ray crystal structure^56^ and Avl9-Arf1 structural prediction. (**B**) *Left,* schematic denoting the subregions of the tripartite DENN domain. *Right,* comparison of the crystal structure of the Rab-GEF DENND1B interacting with Rab35^58^ to the predicted structure of the Avl9-Arf1 interaction. Colors illustrate the subdomain structure. (**C**) Cryo-EM structures of the catalytic subunits of the longin domain GAPs bound to their substrate GTPases, colored by subunit^60,62,64^. (**D**) Comparison of DENN1B, Avl9 prediction, and NRPL2 structures colored by subdomain to illustrate the distinct configuration of the longin domain GAP subunits.

**Figure S6.**
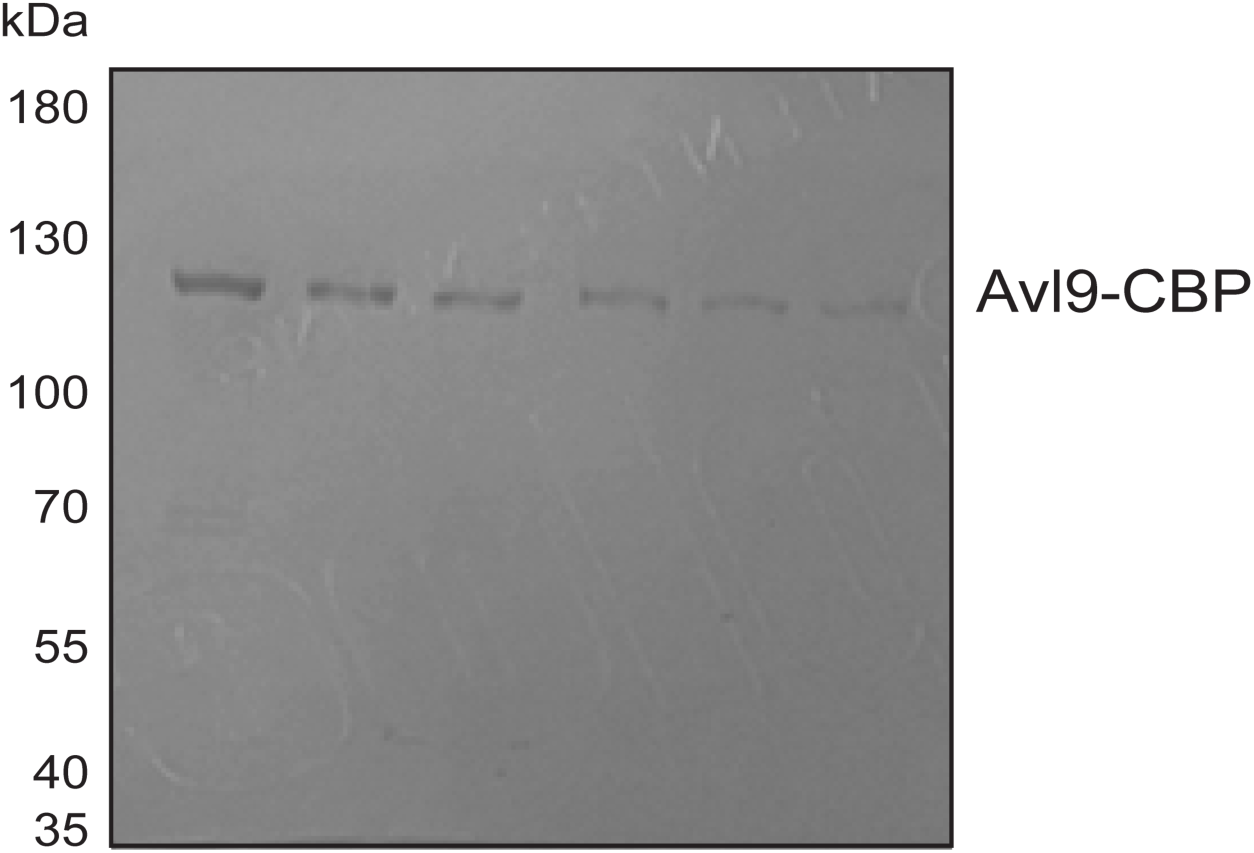
Purification of native Avl9. SDS-PAGE gel stained with Coomassie. Shown are elution fractions from purification of native *S. cerevisiae* Avl9 with fused calmodulin binding peptide (CBP).

**Figure S7.**
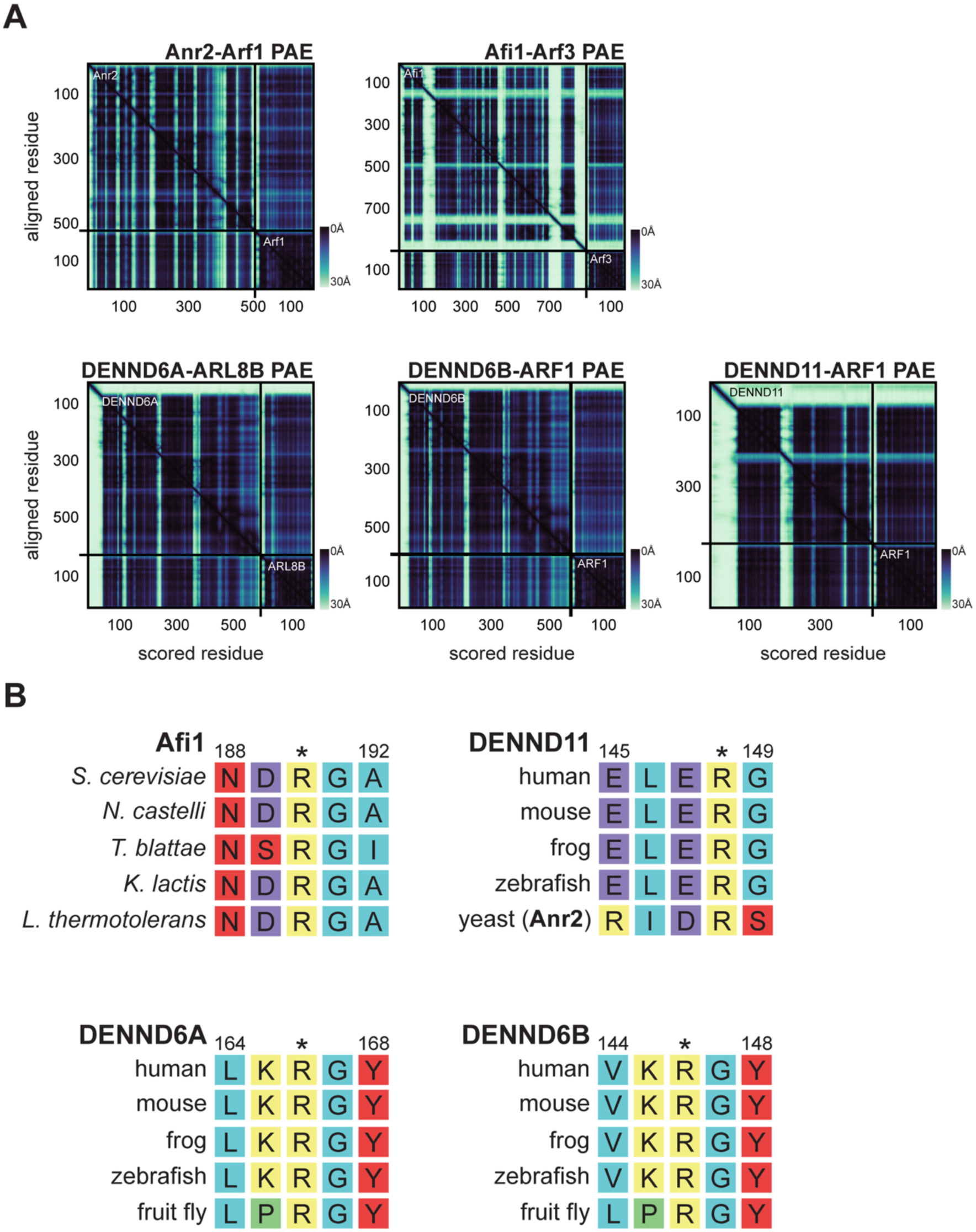
Sequence conservation of predicted arginine finger shared between candidate DENN GAPs. (**A**) PAE plots of candidate DENN GAP-GTPase predictions (**B**) Multiple sequence alignments of candidate DENN GAP proteins with their homologs in other species. Numbers correspond to amino acids of topmost protein. Predicted catalytic arginine, based on structural predictions, shown with asterisk. For context, levels of genome sequence divergence between *S. cerevisiae* and *K. lactis* is similar to that between humans and starfish^97^.

**Figure S8.**
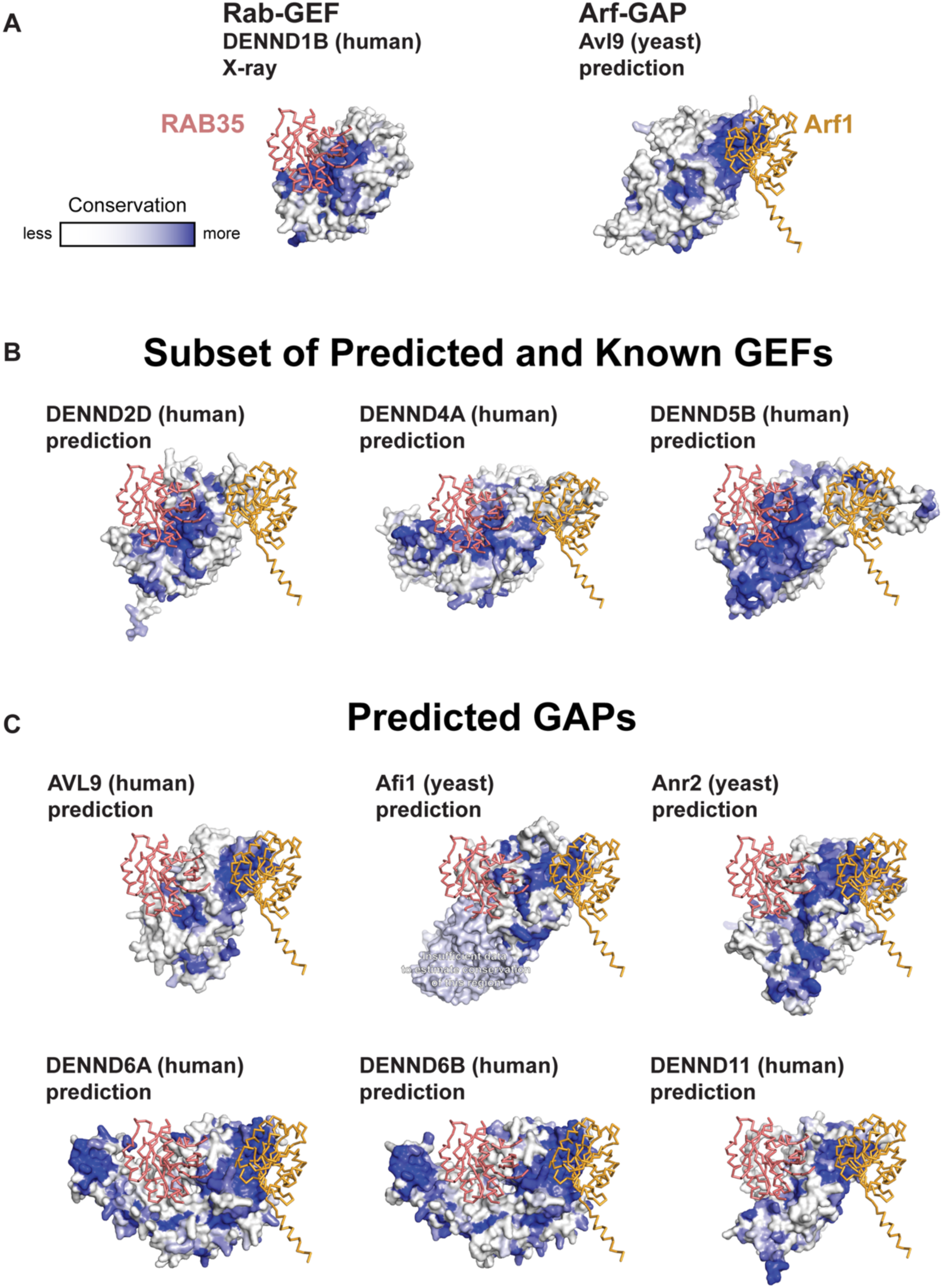
Sequence conservation of “Arf-GAP surface” shared between candidate DENN GAPs. ConSurf analyses of predicted DENN domain structures^98^. (**A**) Rab35 and Arf1 overlaid on each structure based on alignment to known Rab-GEF (DENND1B)^58^ and Arf-GAP (Avl9). (**B, C**) Higher degree of sequence conservation at either Rab35 or Arf1 binding sites indicates likelihood of GEF or GAP function of a given DENN domain protein.

**Figure S9.**
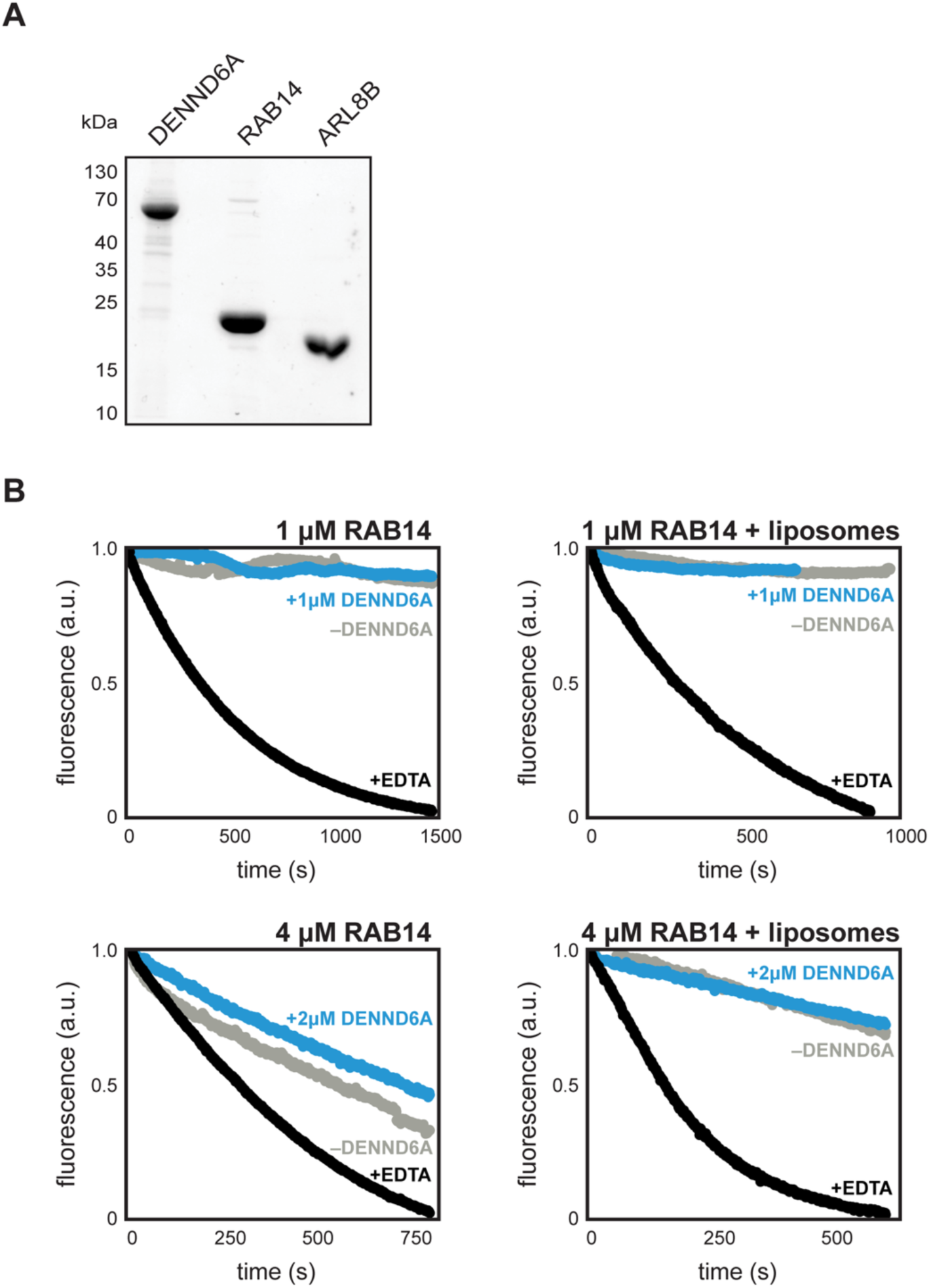
DENND6A *in vitro* GEF assays. (**A**) SDS-PAGE gel stained with Coomassie showing purified DENND6A, mantGDP-RAB14, and ΔN-ARL8B. (**B**) mantGDP fluorescence GEF assay with RAB14-His. Shown are traces for reactions with DENND6A (blue), without DENND6A (mock; gray) and EDTA (positive control; black).

**Table S1.**
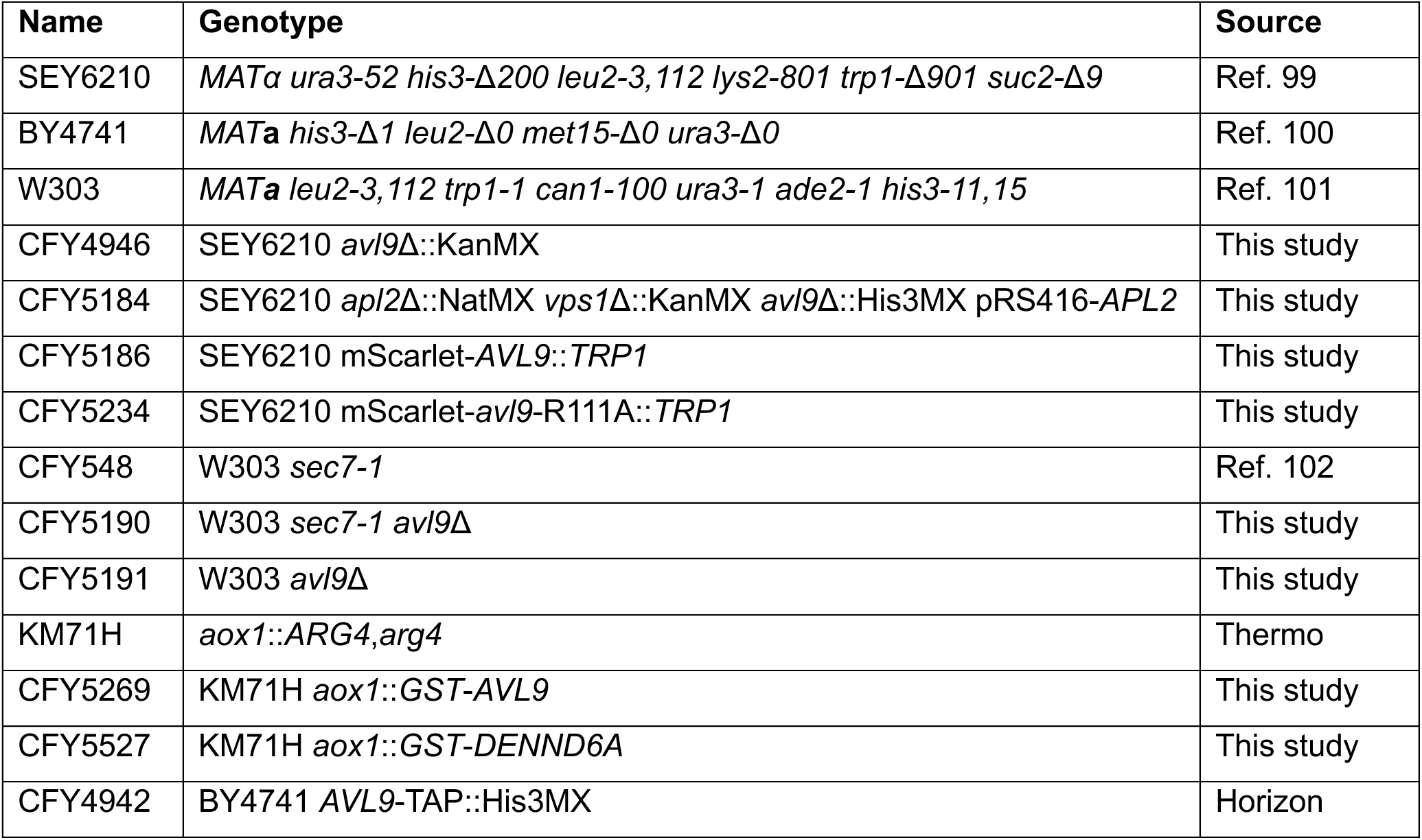
Yeast strains used in this study.

**Table S2.**
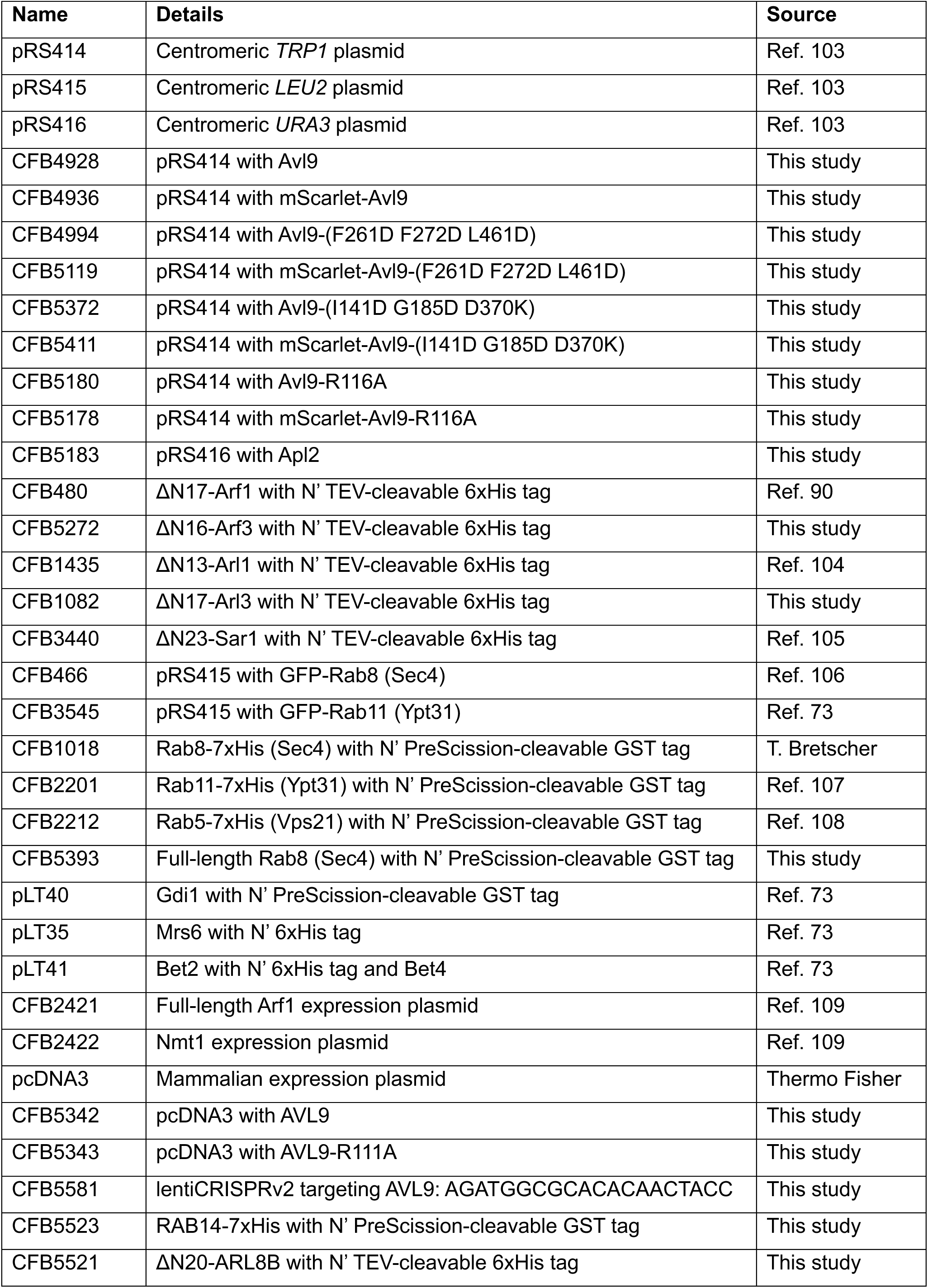
Plasmids used in this study.

**Table S3.** Liposome composition.

| Lipid | mol % |  |
| --- | --- | --- |
|  | Golgi | Ni<sup>2+</sup> Golgi |
| DOPC | 25 | 20 |
| POPC | 6 | 6 |
| DOPE | 7 | 7 |
| POPE | 3 | 3 |
| DOPS | 1 | 1 |
| POPS | 2 | 2 |
| DOPA | 1 | 1 |
| POPA | 2 | 2 |
| Liver PI | 29 | 29 |
| PI(4)P | 1 | 1 |
| CDP-DAG | 2 | 2 |
| PO-DAG | 4 | 4 |
| DO-DAG | 2 | 2 |
| Ceramide (C18) | 5 | 5 |
| Cholesterol | 10 | 10 |
| Ni<sup>2+</sup>-DOGS | 0 | 5 |

## REFERENCES

1. Pfeffer, S. R. Rab GTPases: master regulators that establish the secretory and endocytic pathways. Mol. Biol. Cell 28, 712–715 (2017).

2. Jackson, C. L. & Bouvet, S. Arfs at a Glance. J. Cell Sci. 127, 4103–4109 (2014).

3. Mosaddeghzadeh, N. & Ahmadian, M. R. The RHO Family GTPases: Mechanisms of Regulation and Signaling. Cells 10, 1831 (2021).

4. Mozzarelli, A. M., Simanshu, D. K. & Castel, P. Functional and structural insights into RAS effector proteins. Mol. Cell 84, 2807–2821 (2024).

5. Wennerberg, K., Rossman, K. L. & Der, C. J. The Ras superfamily at a glance. J. Cell Sci. 118, 843–846 (2005).

6. Bos, J. L., Rehmann, H. & Wittinghofer, A. GEFs and GAPs: Critical Elements in the Control of Small G Proteins. Cell 129, 865–877 (2007).

7. Vigil, D., Cherfils, J., Rossman, K. L. & Der, C. J. Ras superfamily GEFs and GAPs: validated and tractable targets for cancer therapy? Nat. Rev. Cancer 10, 842–857 (2010).

8. Donaldson, J. G. & Jackson, C. L. ARF family G proteins and their regulators: roles in membrane transport, development and disease. Nat. Rev. Mol. Cell Biol. 12, 362–375 (2011).

9. Stalder, D. & Antonny, B. Arf GTPase regulation through cascade mechanisms and positive feedback loops. FEBS Lett. 587, 2028–2035 (2013).

10. Chen, P.-W., Gasilina, A., Yadav, M. P. & Randazzo, P. A. Control of cell signaling by Arf GTPases and their regulators: Focus on links to cancer and other GTPase families. Biochim. Biophys. Acta BBA - Mol. Cell Res. 1869, 119171 (2022).

11. Thomas, L. L. & Fromme, J. C. Extensive GTPase crosstalk regulates Golgi trafficking and maturation. Curr. Opin. Cell Biol. 65, 1–7 (2020).

12. Kumari, S. & Mayor, S. ARF1 is directly involved in dynamin-independent endocytosis. Nat. Cell Biol. 10, 30–41 (2008).

13. Enkler, L. et al. Arf1 coordinates fatty acid metabolism and mitochondrial homeostasis. Nat. Cell Biol. 25, 1157–1172 (2023).

14. Ackema, K. B. et al. The small GTPase Arf1 modulates mitochondrial morphology and function. EMBO J. 33, 2659–2675 (2014).

15. Wilfling, F. et al. Arf1/COPI machinery acts directly on lipid droplets and enables their connection to the ER for protein targeting. eLife 3, e01607 (2014).

16. Jumper, J. et al. Highly accurate protein structure prediction with AlphaFold. Nature 596, 583–589 (2021).

17. Evans, R. et al. Protein complex prediction with AlphaFold-Multimer. 2021.10.04.463034 Preprint at 10.1101/2021.10.04.463034 (2022).

18. Humphreys, I. R. et al. Computed structures of core eukaryotic protein complexes. Science 374, eabm4805 (2021).

19. Zhang, J. et al. Predicting protein-protein interactions in the human proteome. Science 0, eadt1630 (2025).

20. Burke, D. F. et al. Towards a structurally resolved human protein interaction network. Nat. Struct. Mol. Biol. 30, 216–225 (2023).

21. Schmid, E. W. & Walter, J. C. Predictomes, a classifier-curated database of AlphaFold-modeled protein-protein interactions. Mol. Cell 85, 1216–1232.e5 (2025).

22. Harsay, E. & Schekman, R. Avl9p, a Member of a Novel Protein Superfamily, Functions in the Late Secretory Pathway. Mol. Biol. Cell 18, 1203–1219 (2007).

23. Wu, Q., Chen, J. D. & Zhou, Z. AVL9 promotes colorectal carcinoma cell migration via regulating EGFR expression. Biol. Proced. Online 24, 1 (2022).

24. Li, D. et al. AVL9 is Upregulated in and Could Be a Predictive Biomarker for Colorectal Cancer. Cancer Manag. Res. 13, 3123–3132 (2021).

25. Zhang, W. et al. Hypoxia-regulated lncRNA CRPAT4 promotes cell migration via regulating AVL9 in clear cell renal cell carcinomas. OncoTargets Ther. 11, 4537–4545 (2018).

26. Linford, A. et al. Rab14 and Its Exchange Factor FAM116 Link Endocytic Recycling and Adherens Junction Stability in Migrating Cells. Dev. Cell 22–540, 952 (2012).

27. Yoshimura, S., Gerondopoulos, A., Linford, A., Rigden, D. J. & Barr, F. A. Family-wide characterization of the DENN domain Rab GDP-GTP exchange factors. J. Cell Biol. 191, 367–381 (2010).

28. Allaire, P. D. et al. The Connecdenn DENN Domain: A GEF for Rab35 Mediating Cargo-Specific Exit from Early Endosomes. Mol. Cell 37, 370–382 (2010).

29. Zhang, D., Iyer, L., He, F. & Aravind, L. Discovery of Novel DENN Proteins: Implications for the Evolution of Eukaryotic Intracellular Membrane Structures and Human Disease. Front. Genet. 3, (2012).

30. Wada, M. et al. Isolation and Characterization of a GDP/GTP Exchange Protein Specific for the Rab3 Subfamily Small G Proteins *. J. Biol. Chem. 272, 3875–3878 (1997).

31. Marat, A. L. & McPherson, P. S. The connecdenn family, Rab35 guanine nucleotide exchange factors interfacing with the clathrin machinery. J. Biol. Chem. 285, 10627–10637 (2010).

32. Yu, X., Breitman, M. & Goldberg, J. A Structure-Based Mechanism for Arf1-Dependent Recruitment of Coatomer to Membranes. Cell 148, 530–542 (2012).

33. Chardin, P. et al. A human exchange factor for ARF contains Sec7- and pleckstrin-homology domains. Nature 384, 481–484 (1996).

34. Yanguas, F., Moscoso-Romero, E. & Valdivieso, M.-H. Ent3 and GGA adaptors facilitate diverse anterograde and retrograde trafficking events to and from the prevacuolar endosome. Sci. Rep. 9, 10747 (2019).

35. Santos, B. & Snyder, M. Sbe2p and Sbe22p, Two Homologous Golgi Proteins Involved in Yeast Cell Wall Formation. Mol. Biol. Cell 11, 435–452 (2000).

36. Shiba, T. et al. Molecular mechanism of membrane recruitment of GGA by ARF in lysosomal protein transport. Nat. Struct. Mol. Biol. 10, 386–393 (2003).

37. Goldberg, J. Structural basis for activation of ARF GTPase: mechanisms of guanine nucleotide exchange and GTP-myristoyl switching. Cell 95, 237–248 (1998).

38. Ahmadian, M. R., Stege, P., Scheffzek, K. & Wittinghofer, A. Confirmation of the arginine-finger hypothesis for the GAP-stimulated GTP-hydrolysis reaction of Ras. Nat. Struct. Biol. 4, 686–689 (1997).

39. Kahn, R. A. et al. Consensus nomenclature for the human ArfGAP domain-containing proteins. J. Cell Biol. 182, 1039–1044 (2008).

40. Marat, A. L., Dokainish, H. & McPherson, P. S. DENN Domain Proteins: Regulators of Rab GTPases. J. Biol. Chem. 286, 13791–13800 (2011).

41. Du, L. L., Collins, R. N. & Novick, P. J. Identification of a Sec4p GTPase-activating protein (GAP) as a novel member of a Rab GAP family. J. Biol. Chem. 273, 3253–3256 (1998).

42. Cuthbert, E. J., Davis, K. K. & Casanova, J. E. Substrate specificities and activities of AZAP family Arf GAPs in vivo. Am. J. Physiol.-Cell Physiol. 294, C263–C270 (2008).

43. Engel, S. R. et al. Saccharomyces Genome Database: advances in genome annotation, expanded biochemical pathways, and other key enhancements. Genetics 229, iyae185 (2025).

44. Manzer, K. M. & Fromme, J. C. The Arf-GAP Age2 localizes to the late-Golgi via a conserved amphipathic helix. Mol. Biol. Cell 34, ar119 (2023).

45. Antonny, B., Huber, I., Paris, S., Chabre, M. & Cassel, D. Activation of ADP-ribosylation Factor 1 GTPase-Activating Protein by Phosphatidylcholine-derived Diacylglycerols *. J. Biol. Chem. 272, 30848–30851 (1997).

46. Yofe, I. et al. One library to make them all: streamlining the creation of yeast libraries via a SWAp-Tag strategy. Nat. Methods 13, 371–378 (2016).

47. Huh, W.-K. et al. Global analysis of protein localization in budding yeast. Nature 425, 686–691 (2003).

48. Tkach, J. M. et al. Dissecting DNA damage response pathways by analysing protein localization and abundance changes during DNA replication stress. Nat. Cell Biol. 14, 966–976 (2012).

49. Poon, P. P., Nothwehr, S. F., Singer, R. A. & Johnston, G. C. The Gcs1 and Age2 ArfGAP proteins provide overlapping essential function for transport from the yeast trans-Golgi network. J. Cell Biol. 155, 1239–1250 (2001).

50. Giaever, G. et al. Functional profiling of the Saccharomyces cerevisiae genome. Nature 418, 387–391 (2002).

51. Poon, P. P. et al. Retrograde transport from the yeast Golgi is mediated by two ARF GAP proteins with overlapping function. EMBO J. 18, 555–564 (1999).

52. Chen, Y. G. & Shields, D. ADP-ribosylation factor-1 stimulates formation of nascent secretory vesicles from the trans-Golgi network of endocrine cells. J. Biol. Chem. 271, 5297–5300 (1996).

53. Beck, R. et al. Membrane curvature induced by Arf1-GTP is essential for vesicle formation. Proc. Natl. Acad. Sci. 105, 11731–11736 (2008).

54. Goud, B., Salminen, A., Walworth, N. C. & Novick, P. J. A GTP-binding protein required for secretion rapidly associates with secretory vesicles and the plasma membrane in yeast. Cell 53, 753–768 (1988).

55. Abramson, J. et al. Accurate structure prediction of biomolecular interactions with AlphaFold 3. Nature 630, 493–500 (2024).

56. Ismail, S. A., Vetter, I. R., Sot, B. & Wittinghofer, A. The Structure of an Arf-ArfGAP Complex Reveals a Ca2+ Regulatory Mechanism. Cell 141, 812–821 (2010).

57. Mandiyan, V., Andreev, J., Schlessinger, J. & Hubbard, S. R. Crystal structure of the ARF-GAP domain and ankyrin repeats of PYK2-associated protein β. EMBO J. 18, 6890–6898 (1999).

58. Wu, X. et al. Insights regarding guanine nucleotide exchange from the structure of a DENN-domain protein complexed with its Rab GTPase substrate. Proc. Natl. Acad. Sci. 108, 18672–18677 (2011).

59. Bar-Peled, L. et al. A Tumor suppressor complex with GAP activity for the Rag GTPases that signal amino acid sufficiency to mTORC1. Science 340, 1100–1106 (2013).

60. Jansen, R. M. et al. Structural basis for FLCN RagC GAP activation in MiT-TFE substrate-selective mTORC1 regulation. Sci. Adv. 8, eadd2926 (2022).

61. Tsun, Z.-Y. et al. The Folliculin Tumor Suppressor Is a GAP for the RagC/D GTPases That Signal Amino Acid Levels to mTORC1. Mol. Cell 52, 495–505 (2013).

62. Su, M.-Y., Fromm, S. A., Remis, J., Toso, D. B. & Hurley, J. H. Structural basis for the ARF GAP activity and specificity of the C9orf72 complex. Nat. Commun. 12, 3786 (2021).

63. Su, M.-Y., Fromm, S. A., Zoncu, R. & Hurley, J. H. Structure of the C9orf72 ARF GAP complex that is haploinsufficient in ALS and FTD. Nature 585, 251–255 (2020).

64. Egri, S. B. et al. Cryo-EM structures of the human GATOR1-Rag-Ragulator complex reveal a spatial-constraint regulated GAP mechanism. Mol. Cell 82, 1836–1849.e5 (2022).

65. Jansen, R. M. & Hurley, J. H. Longin domain GAP complexes in nutrient signalling, membrane traffic and neurodegeneration. FEBS Lett. 597, 750–761 (2023).

66. Kumar, R. et al. DENND6A links Arl8b to a Rab34/RILP/dynein complex, regulating lysosomal positioning and autophagy. Nat. Commun. 15, 919 (2024).

67. Kreider-Letterman, G. et al. ARHGAP17 regulates the spatiotemporal activity of Cdc42 at invadopodia. J. Cell Biol. 222, e202207020 (2022).

68. Gillingham, A. K., Bertram, J., Begum, F. & Munro, S. In vivo identification of GTPase interactors by mitochondrial relocalization and proximity biotinylation. eLife 8, e45916 (2019).

69. Gillingham, A. K., Sinka, R., Torres, I. L., Lilley, K. S. & Munro, S. Toward a Comprehensive Map of the Effectors of Rab GTPases. Dev. Cell 31, 358–373 (2014).

70. Liu, W., Duden, R., Phair, R. D. & Lippincott-Schwartz, J. ArfGAP1 dynamics and its role in COPI coat assembly on Golgi membranes of living cells. J. Cell Biol. 168, 1053–1063 (2005).

71. Rosa-Ferreira, C. & Munro, S. Arl8 and SKIP act together to link lysosomes to kinesin-1. Dev. Cell 21, 1171–1178 (2011).

72. Itzen, A., Rak, A. & Goody, R. S. Sec2 is a Highly Efficient Exchange Factor for the Rab Protein Sec4. J. Mol. Biol. 365, 1359–1367 (2007).

73. Thomas, L. L. & Fromme, J. C. GTPase cross talk regulates TRAPPII activation of Rab11 homologues during vesicle biogenesis. J. Cell Biol. 215, 499–513 (2016).

74. Ortiz, D., Medkova, M., Walch-Solimena, C. & Novick, P. Ypt32 recruits the Sec4p guanine nucleotide exchange factor, Sec2p, to secretory vesicles; evidence for a Rab cascade in yeast. J. Cell Biol. 157, 1005–1016 (2002).

75. Walch-Solimena, C., Collins, R. N. & Novick, P. J. Sec2p Mediates Nucleotide Exchange on Sec4p and Is Involved in Polarized Delivery of Post-Golgi Vesicles. J. Cell Biol. 137, 1495–1509 (1997).

76. Babu, M. et al. Interaction landscape of membrane-protein complexes in Saccharomyces cerevisiae. Nature 489, 585–589 (2012).

77. Ho, Y. et al. Systematic identification of protein complexes in Saccharomyces cerevisiae by mass spectrometry. Nature 415, 180–183 (2002).

78. Michaelis, A. C. et al. The social and structural architecture of the yeast protein interactome. Nature 624, 192–200 (2023).

79. Tarassov, K. et al. An in Vivo Map of the Yeast Protein Interactome. Science 320, 1465–1470 (2008).

80. Horgan, C. P., Hanscom, S. R., Jolly, R. S., Futter, C. E. & McCaffrey, M. W. Rab11-FIP3 links the Rab11 GTPase and cytoplasmic dynein to mediate transport to the endosomal-recycling compartment. J. Cell Sci. 123, 181–191 (2010).

81. Stockhammer, A. et al. ARF1 compartments direct cargo flow via maturation into recycling endosomes. Nat. Cell Biol. 26, 1845–1859 (2024).

82. Casalou, C., Faustino, A. & Barral, D. C. Arf proteins in cancer cell migration. Small GTPases 7, 270–282 (2016).

83. Maritzen, T., Schachtner, H. & Legler, D. F. On the move: endocytic trafficking in cell migration. Cell. Mol. Life Sci. CMLS 72, 2119–2134 (2015).

84. Li, J. H., Trivedi, V. & Diz-Muñoz, A. Understanding the interplay of membrane trafficking, cell surface mechanics, and stem cell differentiation. Semin. Cell Dev. Biol. 133, 123–134 (2023).

85. Tsai, P.-C. et al. Afi1p Functions as an Arf3p Polarization-specific Docking Factor for Development of Polarity *. J. Biol. Chem. 283, 16915–16927 (2008).

86. Currie, E. et al. High confidence proteomic analysis of yeast LDs identifies additional droplet proteins and reveals connections to dolichol synthesis and sterol acetylation [S]. J. Lipid Res. 55, 1465–1477 (2014).

87. Thul, P. J. et al. A subcellular map of the human proteome. Science 356, eaal3321 (2017).

88. Mirdita, M. et al. ColabFold: making protein folding accessible to all. Nat. Methods 19, 679–682 (2022).

89. Luan Ha, V., Thomas, G. M. H., Stauffer, S. & Randazzo, P. A. Preparation of Myristoylated Arf1 and Arf6. in Methods in Enzymology vol. 404 164–174 (Academic Press, 2005).

90. Richardson, B. C., McDonold, C. M. & Fromme, J. C. The Sec7 Arf-GEF Is Recruited to the trans-Golgi Network by Positive Feedback. Dev. Cell 22, 799–810 (2012).

91. Ghaemmaghami, S. et al. Global analysis of protein expression in yeast. Nature 425, 737–741 (2003).

92. Feathers, J. R., Vignogna, R. C. & Fromme, J. C. Structural basis for Rab6 activation by the Ric1-Rgp1 complex. Nat. Commun. 15, 10561 (2024).

93. Klemm, R. W. et al. Segregation of sphingolipids and sterols during formation of secretory vesicles at the trans-Golgi network. J. Cell Biol. 185, 601–612 (2009).

94. Thomas, L. L., van der Vegt, S. A. & Fromme, J. C. A Steric Gating Mechanism Dictates the Substrate Specificity of a Rab-GEF. Dev. Cell 48, 100–114.e9 (2019).

95. Itzen, A., Rak, A. & Goody, R. S. Sec2 is a Highly Efficient Exchange Factor for the Rab Protein Sec4. J. Mol. Biol. 365, 1359–1367 (2007).

96. Suarez-Arnedo, A. et al. An image J plugin for the high throughput image analysis of in vitro scratch wound healing assays. PLoS ONE 15, e0232565 (2020).

97. Shen, X.-X. et al. Tempo and Mode of Genome Evolution in the Budding Yeast Subphylum. Cell 175, 1533–1545.e20 (2018).

98. Yariv, B. et al. Using evolutionary data to make sense of macromolecules with a “face-lifted” ConSurf. Protein Sci. 32, e4582 (2023).

99. Robinson, J. S., Klionsky, D. J., Banta, L. M. & Emr, S. D. Protein Sorting in Saccharomyces cerevisiae: Isolation of Mutants Defective in the Delivery and Processing of Multiple Vacuolar Hydrolases. Mol. Cell. Biol. 8, 4936–4948 (1988).

100. Baker Brachmann, C., et al. Designer deletion strains derived from Saccharomyces cerevisiae S288C: A useful set of strains and plasmids for PCR-mediated gene disruption and other applications. Yeast 14, 115–132 (1998).

101. Replogle, K., Hovland, L. & Rivier, D. H. Designer deletion and prototrophic strains derived from Saccharomyces cerevisiae strain W303-1a. Yeast Chichester Engl. 15, 1141–1149 (1999).

102. Novick, P., Ferro, S. & Schekman, R. Order of events in the yeast secretory pathway. Cell 25, 461–469 (1981).

103. Sikorski, R. S. & Hieter, P. A system of shuttle vectors and yeast host strains designed for efficient manipulation of DNA in Saccharomyces cerevisiae. Genetics 122, 19–27 (1989).

104. Richardson, B. C., Halaby, S. L., Gustafson, M. A. & Fromme, J. C. The Sec7 N-terminal regulatory domains facilitate membrane-proximal activation of the Arf1 GTPase. eLife 5, e12411 (2016).

105. Joiner, A. M. N. & Fromme, J. C. Structural basis for the initiation of COPII vesicle biogenesis. Structure 29, 859–872.e6 (2021).

106. Heger, C. D., Wrann, C. D. & Collins, R. N. Phosphorylation Provides a Negative Mode of Regulation for the Yeast Rab GTPase Sec4p. PLOS ONE 6, e24332 (2011).

107. McDonold, C. M. & Fromme, J. C. Four GTPases differentially regulate the Sec7 Arf-GEF to direct traffic at the trans-golgi network. Dev. Cell 30, 759–767 (2014).

108. Thomas, L. L., Joiner, A. M. N. & Fromme, J. C. The TRAPPIII complex activates the GTPase Ypt1 (Rab1) in the secretory pathway. J. Cell Biol. 217, 283–298 (2018).

109. Schoppe, J. et al. Flexible open conformation of the AP-3 complex explains its role in cargo recruitment at the Golgi. J. Biol. Chem. 297, 101334 (2021).

